# Manipulation of the human tRNA pool reveals distinct tRNA sets that act in cellular proliferation or cell cycle arrest

**DOI:** 10.1101/2020.04.30.070789

**Authors:** Noa Aharon-Hefetz, Idan Frumkin, Yoav Mayshar, Orna Dahan, Yitzhak Pilpel, Roni Rak

## Abstract

Different subsets of the tRNA pool in human are expressed in different cellular conditions. The “proliferation-tRNAs” are induced upon normal and cancerous cell division, while the “differentiation tRNAs” are active in non-dividing, differentiated cells. Here we examine the essentiality of the various tRNAs upon cellular growth and arrest. We established a CRISPR-based editing procedure with sgRNAs that each target a tRNA family. We measured tRNA essentiality for cellular growth and found that most proliferation tRNAs are essential compared to differentiation tRNAs in rapidly growing cell lines. Yet in more slowly dividing lines, the differentiation tRNAs were more essential. In addition, we measured these tRNAs roles upon response to cell cycle arresting signals. Here we detected a more complex behavior with both proliferation-tRNAs and differentiation tRNAs showing various levels of essentiality. These results provide the so-far most comprehensive functional characterization of human tRNAs with intricate roles in various proliferation states.

## Introduction

Cells in multicellular species may typically exist in one of two alternative states, they either proliferate or they are cell cycle-arrested. Differentiated cells are typically less proliferative or they may not divide at all, while proliferation occurs often prior to terminal differentiation, or when differentiation is reversed, predominantly in cancer (Hanahan & Weinberg 2011). The proliferation rate ranges from fast dividing cells (e.g., hematopoietic stem cells), through quiescent cells that divide for replacing dead or injured cells (e.g., fibroblasts, smooth muscle cells, epithelial cells), to cells with little or no proliferation potential (e.g., cardiac muscle tissue) (Rew & Wilson 2000; Ruijtenberg & Van Den Heuvel 2016). Mammalian cells and celllines exit the cell cycle in response to various environmental changes. Quiescence, or the G0-arrest phase is one type of cell cycle arrest state, which is typically invoked in response to nutrient deprivation (Cheung & Rando 2013; Oki et al. 2014; Yao 2014). Senescence is a second pivotal type of cell cycle arrest that is often associated with aging and it is also considered as an anti-cancer mechanism (Collado & Serrano 2010; Pérez-Mancera et al. 2014; Sosa et al. 2014). Tissue homeostasis requires precise and constrict control of these alternative cellular states, and impairment of these regulatory processes may result in degenerative or neoplastic diseases (Besson et al. 2008; Spencer et al. 2013; Yao 2014; Hafner et al. 2019). The regulatory network that controls the proliferation and cell arrest, and the balance between them have been heavily investigated, yet predominantly at the transcription level since data is mostly available at the RNA level (Bar-joseph et al. 2008; Nagano et al. 2016; Hafner et al. 2017; Hernandez-Segura et al. 2017; Casella et al. 2019). Protein translation on the other hand, specifically translation elongation, though studied extensively too (Aviner et al. 2015; Patil et al. 2012; Rapino et al. 2018; Bludau & Aebersold 2020; Knight et al. 2020) remain less characterized in these systems.

tRNAs are a key molecular entity that converts the transcriptome into the proteome. The composition and abundance of the cellular tRNA pool is coordinated to match the codon demand of the transcriptome, which enables optimization of protein synthesis (Dos Reis et al. 2004; Gingold et al. 2012; Gardin et al. 2014; Hanson & Coller 2017; Frumkin et al. 2018). The efficiency of translation is often attributed to the mutual adaptation between supply – the abundance of each tRNA family in the cell, and the demand – the mRNA content of the transcriptome and in particular the extent of usage of each of the 61 types of codons, as demanded by the expressed transcripts. Efficient protein translation depends on supply-to-demand adaptation and it is attained when the highly demanded codons are matched by abundantly available corresponding tRNAs (Presnyak et al. 2015; Hanson & Coller 2017; Rak et al. 2018). Indeed, expression level of tRNAs was shown in recent years not to be constant and to be subject to extensive regulation that in part matches supply to demand (Dittmar et al. 2006; Kirchner & Ignatova 2014; Pan 2018; Rak et al. 2018; Hernandez-Alias et al. 2020). For example, cancerous cells show massive changes in expression of the tRNA pool (Pavon-Eternod et al. 2009; Gingold et al. 2014; Zhang et al. 2018; Santos et al. 2019; Hernandez-Alias et al. 2020). Metastatic cells show typical changes too (Goodarzi et al. 2016).

In particular, it was demonstrated that proliferating and differentiated cells differentially express distinct sets of tRNAs, which tend to match the variable codon usage demand in these conditions (Gingold et al. 2014). In examination of diverse proliferative (normal or cancerous) cells, along with differentiated and arrested cells, we previously defined two sets of human tRNAs, the “proliferation-tRNAs” that are induced in proliferating cells and repressed in differentiated and arrested cells, and the “differentiation-tRNAs” that largely manifest the opposing dynamics. Together, tRNAs from the two sets make up close to half of the human tRNA pool, the rest are tRNAs that show no consistent dynamics of expression across these conditions.

Yet, the correlation between the state of the tRNA pool and the proliferative or arrested cellular state does not reveal causal effects. Are the proliferation-induced tRNAs indeed more essential during cellular proliferation than the differentiation/arrest – induced tRNAs? And which tRNAs are needed during and following the response to cell arresting signals?

To elucidate the functional essentiality of the proliferation-associated, and differentiation-associated tRNAs in diverse cellular states we edited various human tRNA genes using CRISPR-iCas9. We succeeded to systematically CRISPR-target a significant portion of the human tRNA gene families in human cell lines with a single sgRNA per tRNA family. This resulted in a set of cellular clones, on the background of several cell lines, each have a perturbed expression of one tRNA family. We then assess in a pooled competition fashion the relative essentiality of each tRNA family in several cell lines that together span a range of cellular proliferation levels. By and large, the previously defined proliferation-tRNAs were found, as a group to be more essential than the differentiation tRNAs in most highly proliferative cell lines, and less so in slowly proliferating cells. Yet, the essentiality of tRNAs under cell arresting conditions was more complex with members from each tRNA group showing differential essentiality. Our results thus reveal the distinct role of various tRNAs in cellular proliferation and cell cycle arrest.

## Results

### Designing a sgRNA library that targets human tRNA gene families

In this study, we aimed at perturbing the expression of diverse human tRNA genes and to then examine the effects on various cellular phenotypes. tRNA genes appear as isodecoder gene families, i.e. sets of tRNA genes that share the same anti-codon identity. In the human genome, tRNA families can consist of up to dozens of members, with diverse degrees of sequence similarities in the tRNA molecule outside of the anti-codon itself. This situation poses a major challenge in targeting the members of each tRNA family with CRISPR-based genomic editing. In designing the sequences of the sgRNAs to edit each tRNA gene family we attempted to target as many as possible members of the family. In our library design we used a single sgRNA per family, selecting for the one that maximizes coverage across family members. Yet, due to sequence divergence among family members, and additional constraints on sgRNA design, in some families we could at best target a portion of the family members with full complementarity to the sgRNA sequence (Fig. 1A and 1B). Choosing a sgRNA that best fits most of the tRNA family members, thus resulted in targeting of the most conserved sequence elements along the tRNA molecule (Fig 1C), and in targeting the most conserved members from each tRNA family. Inevitably though in many tRNA families some members had mismatches relative to their sgRNA sequence, which suggest that these tRNA genes were not targeted by the single sgRNA. Indeed, by deep-sequencing of the tRNA pool in WT HeLa cells, we found that the members that were fully matched to their respective sgRNA in each family tended to be significantly more highly expressed than those that due to sequence divergence probably evade targeting (Fig 1D, Wilcoxon rank sum test, p < 10^−4^). This result is in line with classical observations that consistently show that the rate of evolution of tRNA genes, i.e., their sequence conservation, correlates with expression level (Thornlow et al. 2018). The practical and desired implication of this correlation between expression level and conservation (and hence targetability in this experiment) means that although for some tRNA families we targeted only a portion of the members, we often targeted the ones that are more highly expressed, i.e. those members that might contribute more to the functional tRNA pool.

**Figure 1.**
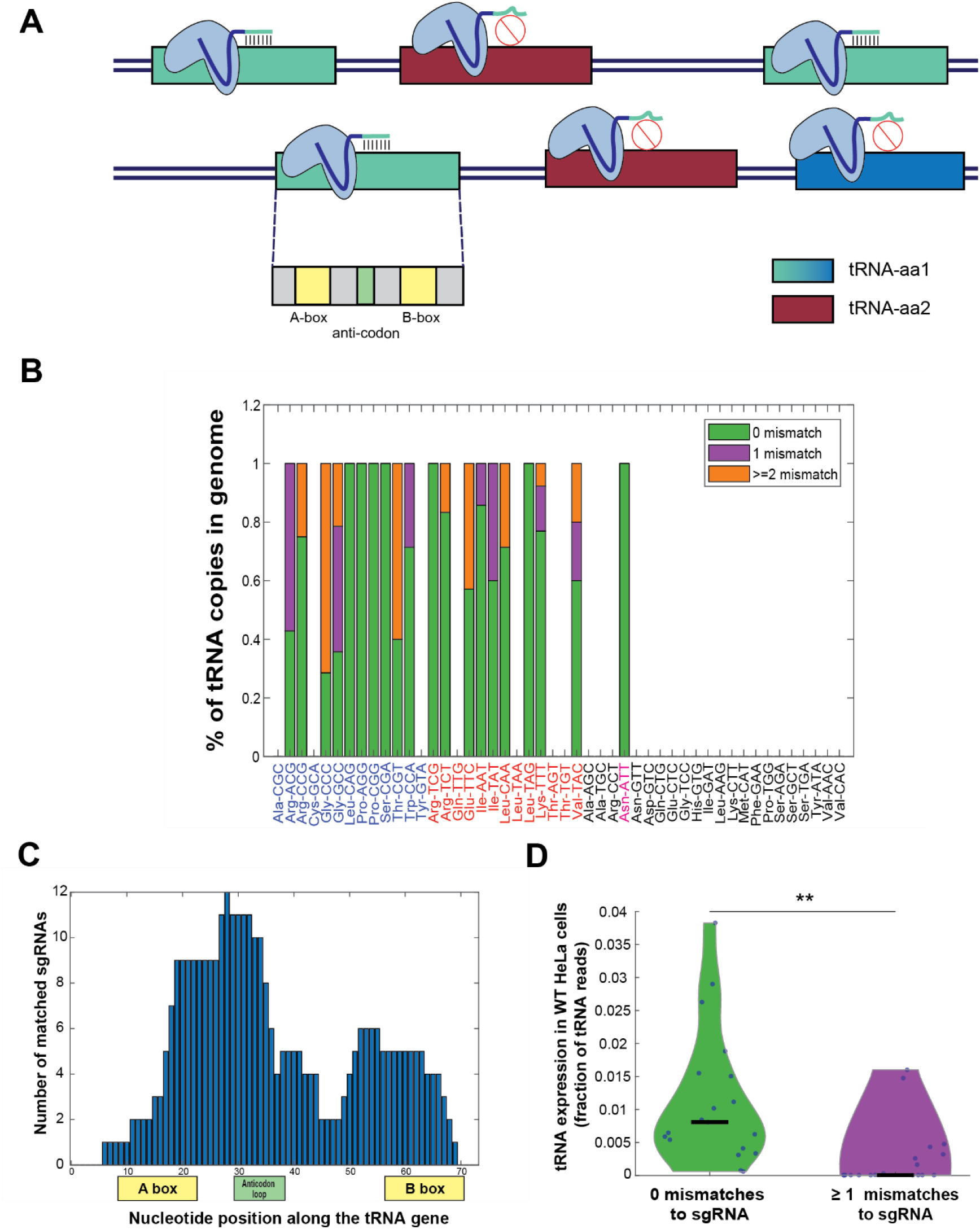
sgRNA library design for genomic editing of human tRNA genes. A| A schema illustrating the sgRNA design for tRNA targeting. The hypothetical tRNA-aa1 family (blue tRNA genes) is targeted by the light-blue sgRNA. 3 out of the 4 tRNA genes (light blue tRNA genes) are fully match to the sgRNA sequence, while the fourth gene has sequence dissimilarities (dark blue tRNA gene), thus predicted not to be targeted by the sgRNA. The hypothetical tRNA-aa2 family (bordeaux tRNA genes) is not predicted to be targeted by the light-blue sgRNA, due to lack of complementarity between the sequences. In addition, the sgRNAs are designed to target functional sequence regions along the tRNA gene to maximize the manipulation effect on the targeted tRNA. B| A bar plot representing the sequence similarity of the tRNA genes to the corresponding sgRNA. Each bar denotes a CRISPR-targeted tRNA family, overall, 19 out of the 46 tRNA families in the human genome and one pseudo tRNA family (Asn-ATT) were targeted. The y-axis denotes the fraction of CRISPR-targeted tRNA genes for each tRNA family (considering only tRNA genes with tRNA score > 50, except of the pseudogene AsnATT tRNA family which consists of two genes with tRNA score <50 (colored in pink)) (Lowe & Chan 2016). The colors in the bars describe the variety of sequence similarity of the tRNA family to the sgRNA sequence (full match in green, 1 mismatch in purple and 2 or more mismatches in orange). The tRNA identity is colored according to the differentiation (blue) / proliferation (red)/ others (black) classification. C| A histogram of the location of the sgRNA sequences along the tRNA genes. The x-axis depicts the nucleotide position along the tRNA, with the A box, B box, and the anticodon loop marked. The y-axis depicts the number of sgRNAs that are complementary to each nucleotide. D| A violin plot describing the distribution of expression levels, in HeLa cells, of perfectly CRISPR-targeted tRNA genes, i.e. that are fully matched to their corresponding sgRNA and non-perfectly CRISPR-targeted tRNA genes, i.e. that have mismatched sequence relative to their corresponding sgRNA. For the fully matched tRNAs and the mismatched tRNAs, the expression of each tRNA family in HeLa cells was calculated by the sum expression of the relevant tRNA copy. The distributions are significantly different, -Wilcoxon rank sum test, p <10^−4^.

Another challenge in applying a CRISPR-based genomic editing of tRNAs is their non-coding nature. When exploiting CRISPR-Cas9 based editing on protein coding genes, most of the disruptive effects result from out-of-frame Indel mutations upon repair. Yet, in non-coding genes, the effect of Indels on functionality are less trivial due to lack of a reading frame (Ho et al. 2015). However, the tRNA gene includes sequence elements that are critical for the functionality, such as the anticodon loop or the internal promoter. During the sgRNA library design, we attempted to direct the sgRNAs to the functional sequence elements of the tRNA (Fig 1A and S1A), so that potentially near-by indels would be maximally perturbing. A point in favor of our approach is that tRNAs are among the shortest RNAs in the human transcriptome, hence Indels, of even a few bases constitute a significant portion of the molecule and will thus disrupt the secondary structure of the molecule, hence reducing functionality of the mature tRNA. A last challenge in this sgRNA library design is the potential for off-target effects between different tRNA families, or non-tRNA genes. Yet we succeeded to minimize potential off-targeting by a design of the single sgRNAs for each family that has at most a minimal complementarity to off-target tRNAs or non-tRNA genes (Fig S1B).

In total we have targeted 19 out of 46 families of human tRNA genes, of which 9 that constitute the “proliferation-tRNAs”, 10 that constitute the “differentiation-tRNAs” (Gingold et al. 2014). In addition, we targeted one pseudo-tRNA family, AsnATT.

### Genomic editing of proliferation tRNAs results in negative selection and a global change of the tRNA pool in HeLa cells

The “proliferation tRNAs” were previously defined as tRNAs that are induced in cancer and other proliferating cells in comparison to other “differentiation tRNAs” that are induced in differentiated tissues (Gingold et al. 2014). To test whether proliferating cells are affected differently by the targeting of these two sets of tRNAs and to evaluate the effect of genomic editing on the tRNA expression and maturation process, we begin by the targeting of a selection of four tRNAs in HeLa cells – two proliferation tRNAs, LeuTAG and ArgTCG, and two differentiation tRNAs, ProCGG and SerCGA. Since all tRNA isodecoder genes of these four tRNA families are perfectly complementary to their respective sgRNA sequences, they are predicted to be fully targeted by the CRISPR system (Fig 1B). We transduced HeLa cells expressing an inducible Cas9 (i.e., iCas9) with each of the four sgRNAs, separately and independently (Fig 2A). Following antibiotic selection, we induced the iCas9 gene by adding Doxycycline to the cell’s media for 12 days. Then, we performed RNA sequencing of the mature tRNAs in each cell population in four time points along the iCas9 induction. Lastly, we estimated the fold-change in expression of the CRISPR-targeted tRNAs in treated cells relative to the WT cells (see Martials and Methods).

**Figure 2.**
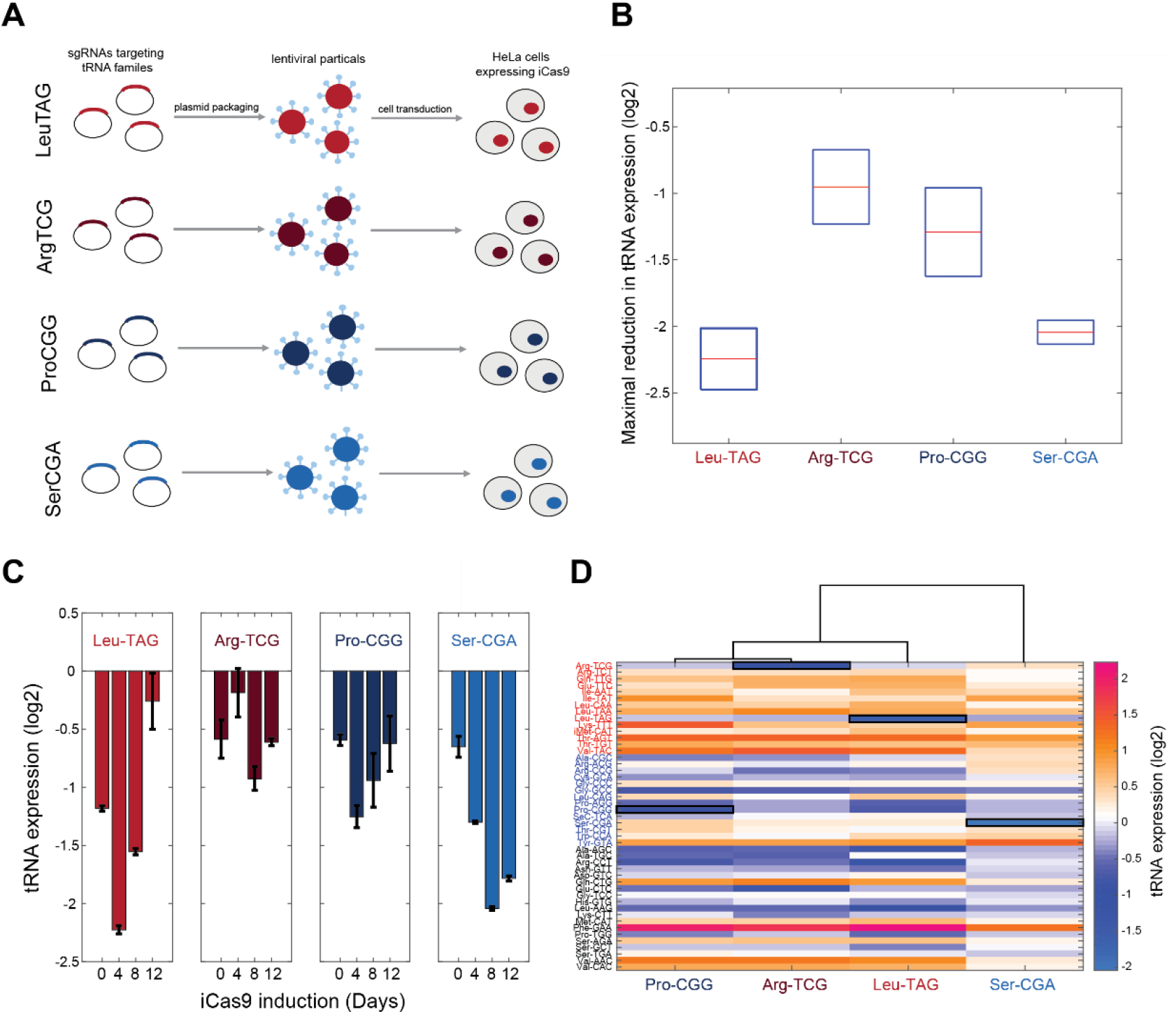
Genomic editing of proliferation-demanded tRNAs exerts a negative selection and a global change in the cellular tRNA pool in HeLa cells. A| A conceptual illustration of the individual tRNA targeting in HeLa cells. Four tRNAs are targeted each by a specific sgRNA. Each sgRNA plasmid was packed in lentiviral particles separately and independently, which then transduced into HeLa cells expressing iCas9. B| A box plot representing the maximal reduction in expression of each CRISPR-targeted tRNA. The y-axis depicts the maximal fold-change (log2) in the tRNA expression of the CRISPR-targeted tRNA in the treated cells relative to WT cells (see Materials and Methods). The x-axis denotes the different tRNA families. C| A bar plot describing the tRNA expression dynamics along the iCas9 induction for each CRISPR-targeted tRNA. The fold change in tRNA expression was calculated as described above. Each bar represents a time point during the iCas9 induction. The two left plots describe the proliferation edited tRNA samples and the two left plots describe the differentiation edited tRNA samples. D| A heat map representing the differential expression of the cellular tRNA pool in CRISPR-targeted tRNA cells. Each column represents a CRISPR-targeted tRNA sample, and each row represents a tRNA isodecoder, grouped by their type (proliferation-red/differentiation-blue/other-black). The color code depicts the fold-change (log2) in tRNA expression in the CRISPR-targeted tRNA sample at day8 of the iCas9 induction relative to the WT sample at the same day. The expression level of the CRISPR-targeted tRNA in each sample is marked in black square. The dendrogram represents the hierarchical clustering of the different CRISPR-targeted tRNA cells based on changes in tRNA expression profile.

The expression level of the CRISPR-targeted tRNAs reduced up to 2-fold compared to WT HeLa cells, an indication for the effectiveness of the genomic editing (Figure 2B). Yet, we noticed that for each CRISPR-targeted tRNA, the maximum reduction in the expression level was reached at a different day following the iCas9 induction. In particular, LeuTAG and ProCGG were maximally repressed already at day 4, while only at day 8 ArgTCG and SerCGA reached to the lowest expression level. We then tested the expression pattern of each CRISPR-targeted tRNA throughout the iCas9 induction for each treated population. For LeuTAG, ProCGG and SerCGA, we found a similar pattern, i.e. a decrease in expression level, followed by a gradual recovery to basal levels (Figure 2C). However, even at day 12 of the iCas9 induction, the expression level of SerCGA remain low (Fig 2C. Day8 vs Day12, t-test, p = ns). These differences between the tRNAs suggest that cells with CRISPR-edited SerCGA genes have a higher fitness relative to WT cells compare to CRISPR-edited LeuTAG and ProCGG genes. To test this hypothesis, for each targeted tRNA, we computed the ratio between the number of reads corresponding to the CRISPR-edited tRNAs and the total tRNA reads, both at the genomic level (by DNA sequencing (Fig S2A)) and at the RNA level (from the RNA sequencing (Fig S2B)). At both levels, we found that the dynamics of the targeted tRNA fraction along the iCas9 induction reflected the expression level. An increase in the CRISPR-edited tRNA reads in the first 4-8 days of the iCas9 induction is followed by an increase of the intact tRNA reads on the expense of the CRISPR-edited tRNA reads (Fig S2). Together, the observed dynamics suggest that most CRISPR editing occurs in the first four to eight days (Yuen et al. 2017), and it is then followed by a competition between cells with various types and extents of editing. The cell competition results in a decline in the edited form of the CRISPR-targeted tRNAs that reflects selection against edited tRNA cells. The negative selection appears to be most pronounced for LeuTAG and ProCGG edited cells suggesting that these tRNAs are essential in proliferative Hela cells. Conversely, cells with CRISPR-edited SerCGA genes had only a minor disadvantage and mainly continued to propagate in the population along with their un-edited counterparts. Moreover, these results propose that ProCGG tRNA, despite its original designation as a differentiation tRNA, might have an essential role in proliferation of HeLa cells.

In addition to changes in the expression levels of the CRISPR-targeted tRNA itself, the tRNA pool in cells may feature a more complex response following the reduction of each individually targeted tRNAs. Such dynamics would be reflected in expression changes of other tRNAs that were not CRISPR-targeted directly. We, therefore, monitored, in HeLa cells, the change in the expression levels of all tRNAs in each of the four individually manipulated tRNA populations throughout iCas9 induction. We observed that the tRNA pool does respond to genomic editing of individual tRNAs, either by induction or repression of tRNAs, by factors as high as 2 to 4 (Fig 2D). Further, the tRNA pool responded similarly in the CRISPR-targeting of the three tRNAs that proved to be the most essential in the above experiment, namely ProCGG, ArgTCG and LeuTAG (Fig 2C and S2). In contrast, the response to the targeting of SerCGA, which is relatively less essential in HeLa cells, was mild also at the tRNA pool level and it resembled the pool of WT cells (Fig 2D). When examining the type of tRNAs which are differentially expressed in ProCGG, ArgTCG and LeuTAG-targeted cells, we observed that most of the up-regulated tRNAs belong to the proliferation tRNA group, while most of the down-regulated tRNAs belong to the differentiation tRNA group (Fig 2D). The other tRNAs, those that do not belong to the proliferation or differentiation tRNA sets, showed a mixed pattern that was characterized with either up or down expression regulation (Fig 2D). These results suggest that the tRNA pool in HeLa cells is responsive to the reduction of essential tRNAs. In particular, proliferation tRNAs are preferentially up-regulated, whereas differentiation tRNAs are mostly down-regulated following expression manipulation of essential tRNAs.

### Proliferation tRNAs are more essential than differentiation tRNAs for HeLa cellular growth

As we validated that CRISPR-iCas9 is a suitable system to perturb the tRNA expression level, we conducted a CRISPR-targeted tRNA competition experiment among our designed library of 20 sgRNAs. We transduced HeLa-iCas9 cells with the sgRNA library in a way, in which each cell in the population expresses a single sgRNA type, and induced the iCas9 gene for 14 days (Fig 3A). To evaluate the growth dynamics of each of the CRISPR-targeted tRNA variants among the competing cells in the population, we deep-sequenced the genomic region encoding for the sgRNAs in five time points during the iCas9 induction. Then, we estimated the relative fitness of each CRISPR-targeted tRNA variant by calculating the fold change of the sgRNA frequency in each time point relative to day 0 (before iCas9 induction) (Fig 3A, see Materials and Methods). Beginning from day 7 of the competition, we observed a strong difference in relative sgRNA frequency between the CRISPR-targeted tRNA groups, with some CRISPR mutants declining in frequency by 2 orders of magnitude already at day 7, and by 3 orders of magnitude at day 14 relative to day zero (Fig 3B). Many of the targeted proliferation tRNA variants showed a sharp decline in frequency in the pooled population, while many of the targeted differentiation tRNA variants showed relatively mild change in frequency (Fig 3B). This is a clear indication that on average, as a group, the proliferation tRNAs are more essential for HeLa cells. Assuming an unperturbed doubling of about one day for this cell line, a decline in frequency of some of the proliferation tRNAs by a factor of ~4,000 compare to the total targeted population, obtained over 14 days, indicates almost complete arrest of cell doublings or cell death upon iCas9 activation. CRISPR-targeting of the pseudo tRNA AsnATT, a very lowly expressed tRNA in HeLa cells, elevated the relative frequency of the cells that carried its sgRNA, indicating at most small contribution to fitness (Fig 3B).

**Figure 3.**
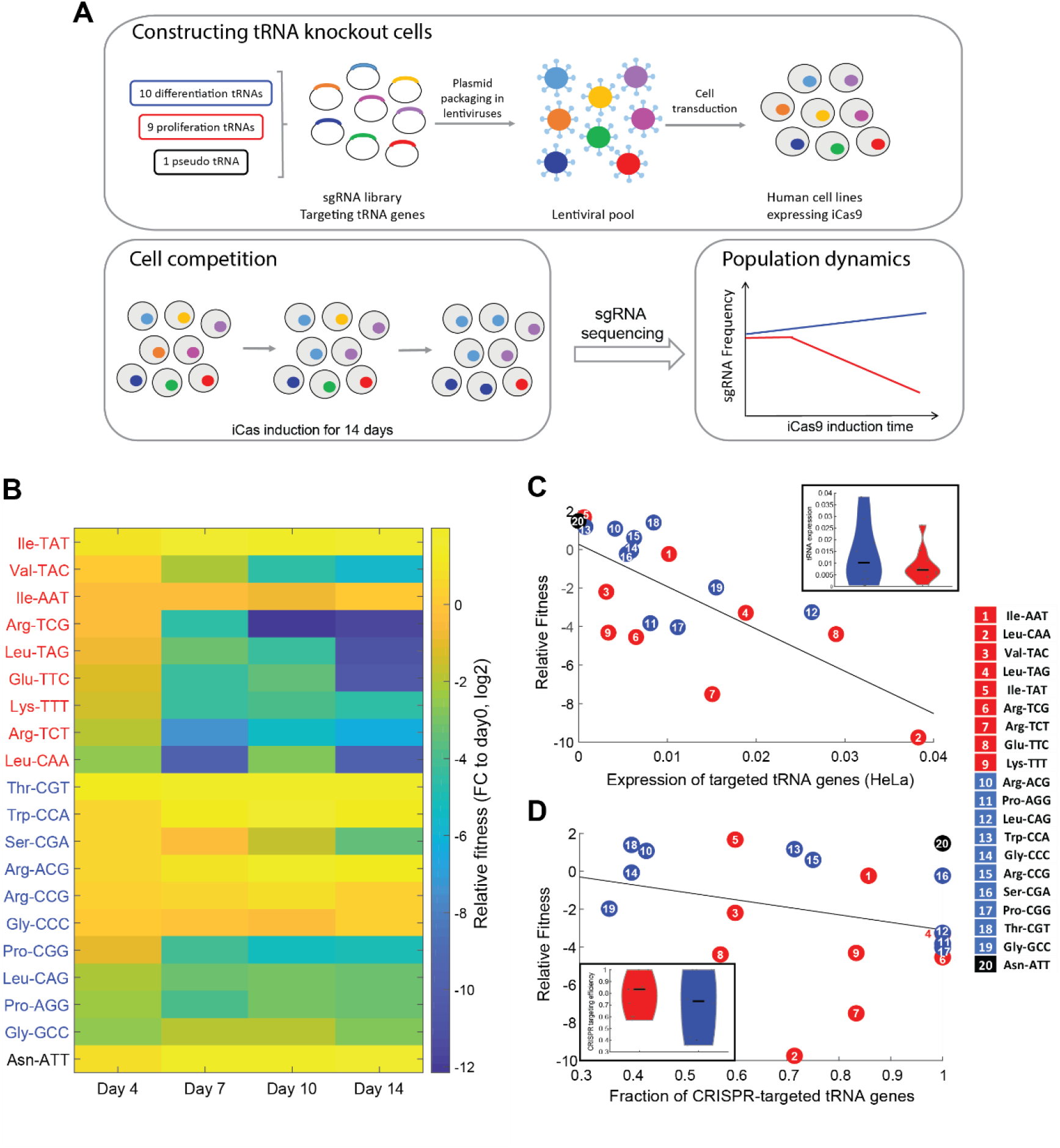
Proliferation tRNA are essential for cellular growth in HeLa cells. A| The experimental design. We designed a CRISPR-sgRNA library, in which each sgRNA targets specific tRNA gene family. Following cloning of the sgRNAs into a lenti-plasmid, we produced a lenti-viral pool that contained the entire sgRNA pool. Then, we transduced human cell lines (HeLa, WI38 fast and WI38 slow) expressing an inducible Cas9 with the lenti-viral sgRNA pool. We performed a CRISPR-edited tRNA cell competition by induction of the iCas9 in parallel to antibiotic selection (two biological repeats for each cell line). The iCas9 induction continued for 14 days, while we sampled the heterogonous population every 3-4 days. Lastly, we deep-sequenced the sgRNAs in each sample, to evaluate the growth dynamics of the targeted-tRNA cells. B| A heat map representation of the relative fitness of CRISPR-targeted tRNA variants in HeLa cells. Each row represents a CRISPR-targeted tRNA variant. The tRNAs (amino acid and anti-codon) are colored according to the tRNA classification: proliferation tRNAs in red and differentiation tRNAs in blue. The pseudo tRNA is colored in black. Each column represents a time point during the iCas9 induction. The color code depicts a proxy of each row’s relative fitness – fold-change (log2) of the sgRNA read frequency in each time point relative to the sgRNA read frequency in day 0 of the iCas9 induction (see Materials and Methods). The values were averaged over two biological repeats. C| A scatter plot comparing the expression of the CRISPR-targeted tRNAs in WT HeLa cells and the fitness of the CRISPR-targeted tRNA cells in HeLa cell line (Pearson correlation, r = −0.71, p < 10^−3^). The x-axis denotes the expression of the CRISPR-targeted tRNA familys (each is summed by the expression of the tRNA isodecoder genes that perfectly match their corresponding sgRNA sequence) in WT HeLa cells. The y-axis denotes the relative fitness of the CRISPR-targeted tRNA cells in day 7 of the iCas9 induction. Each dot is a CRISPR-targeted tRNA family. The colors and numbers denote the tRNA group (proliferation-red/ differentiation-blue/pseudo-black). A Violin plot representing the difference in expression level between the targeted proliferation tRNAs and the targeted differentiation tRNAs in WT HeLa cells is presented as a subset. Each dot depicts a single tRNA family. The red violin stands for the proliferation tRNA families, while the blue violin stands for the differentiation tRNA families. (Wilcoxon rank-sum test, p = ns) D| A scatter plot comparing the fraction of CRISPR-targeted tRNA genes for each sgRNA and the relative fitness of the CRISPR-targeted tRNA cells in HeLa cell line (Pearson correlation, (without the pseudo tRNA AsnATT), r = −0.38, p = ns). The x-axis denotes the ratio of the number of fully complemented tRNA isodecoder genes to their corresponding sgRNA to the total number of tRNA isodecoder genes. The y-axis denotes the relative fitness of the CRISPR-targeted tRNA cells in day 7 of the iCas9 induction. The colors and numbers represent the CRISPR-targeted tRNAs classified to proliferation-red/ differentiation-blue/pseudo-black. A violin plot representing the difference in the fraction of CRISPR-targeted tRNA genes between proliferation and differentiation tRNAs in HeLa cells is presented as a subset. Each dot depicts a single tRNA family. The red violin stands for the proliferation tRNA families, while the blue violin stands for the differentiation tRNA families. (Wilcoxon rank-sum test, p = ns).

We were further interested in revealing whether the expression levels of each of the tRNAs by the cells can explain the effect of their CRISPR-targeting on the fitness, and if such expression levels differ between proliferation and differentiation tRNAs in this cell line. For that, we examined the correlation between the expression levels of each of the CRISPR-targeted tRNAs in WT HeLa cells to the fitness of their edited tRNA variants. We found a negative correlation between the tRNA expression in WT cells and the relative fitness of the CRISPR-targeted tRNA variants, indicating higher essentiality of highly expressed tRNA (Fig 3C, Pearson correlation, r = −0.71, p < 10^−3^). This is in line with classical observations made in evolutionary studies showing that highly expressed genes tend to be more essential than lowly expressed ones (Krylov et al. 2003). We next wanted to examine if the higher essentiality of the proliferation tRNAs could be reduced to their being more highly expressed than the differentiation tRNAs in HeLa cells. Yet we found no significant difference in distribution of expression level between the proliferation and differentiation tRNAs in WT HeLa cells (Fig 3C-violin plot, Wilcoxon rank-sum test, p = ns), indicating that the higher essentiality of the proliferation tRNAs (Fig 3C and Fig S3, Wilcoxon rank-sum test, p < 0.05) cannot be explained by mere expression level difference.

Next, we examined the possibility that the difference in essentiality of the two tRNA sets can be trivialized by a difference in the fraction of tRNA genes that are predicted to be targeted by their sgRNA (i.e the fraction of gene family members that fully match the sgRNA). This concern could have its basis in the observed, yet not significant, negative correlation between the fitness reduction and the fraction of CRISPR-targeted tRNA genes (Fig 3D, Pearson correlation (without the pseudo tRNA AsnATT), r = −0.38, p = ns). Reassuringly, the proportion of targeted family members is similar among the proliferation and differentiation tRNA families (Fig 3D-violin plot, Wilcoxon rank-sum test, p = ns) and we thus exclude this factor too as the explanation for the difference in essentiality of the two tRNA sets.

### The Response to CRISPR-targeting of tRNAs is dependent on the cell line origin and the growth rate

We next moved to examine the essentiality of the various tRNAs in more slow-growing cell lines. We looked for at least two human cell lines of similar origin that yet manifest different growth rates. We chose two fibroblasts cell lines that were both derived from the same original fibroblast cell line, WI38, in a serial passaging process (Milyavsky et al. 2003). An early and late time point along the serial passaging process yielded respectively the “WI38 slow” cell line and the “WI38 fast” cell line, whose doubling times are around ~72 hours and ~ 24h, respectively (compared to HeLa’s ~20 hour).

We compared the relative fitness of different CRISPR-targeted tRNA variants in HeLa and these two additional cell lines. Overall, tRNAs tended to show similar relative fitness scores in all cell lines (Fig 4. Person correlation, HeLa vs WI38 Slow: r = 0.54, p < 0.05; WI38 Fast vs WI38 Slow: r = 0.81, p < 10^−5^; HeLa vs WI38 Fast: r = 0.76, p < 10^−5^). Yet, a more detailed comparison revealed a difference between the cell lines. Comparing tRNA essentiality in WI38 slow cells and HeLa cells, i.e. the slowest to the fastest cell lines in this collection, revealed a marked difference in tRNA essentiality. While HeLa cells were more sensitive to CRISPR-targeting of most of the proliferation tRNAs but only mildly sensitive to differentiation tRNA targeting, WI38 slow cells showed enhanced sensitivity to CRISPR-targeting of differentiation tRNAs, and much lower sensitivity to targeting of proliferation tRNAs (Fig 4A). When comparing WI38 slow to WI38 fast cells, we observed that most tRNA targeting affected the cell’s fitness similarly in the two cell lines, yet, three CRISPR-targeted tRNA variants differed in fitness. Two of the three are proliferation tRNAs that showed lower fitness in WI38 fast cells than in WI38 slow cells, and a third is a differentiation tRNA that was more essential in the WI38 slow cells (Fig 4B). We also observed higher sensitivity of WI38 fast cells to differentiation tRNA editing compared to HeLa cells, although the difference was less pronounced than the effect of differentiation tRNA editing in WI38 slow (Fig 4C). These results indicate that tRNA essentiality depends both on cell origin and on the proliferation status of the cell. In particular, the more proliferative cells show a higher essentiality of the proliferation tRNAs, while slower cell lines’ fitness depend more on differentiation tRNAs. Cellular origin appears relevant too – although the WI38 slow and fast cells differ markedly in doubling time, their essentiality profile is almost identical across most tRNAs.

**Figure 4.**
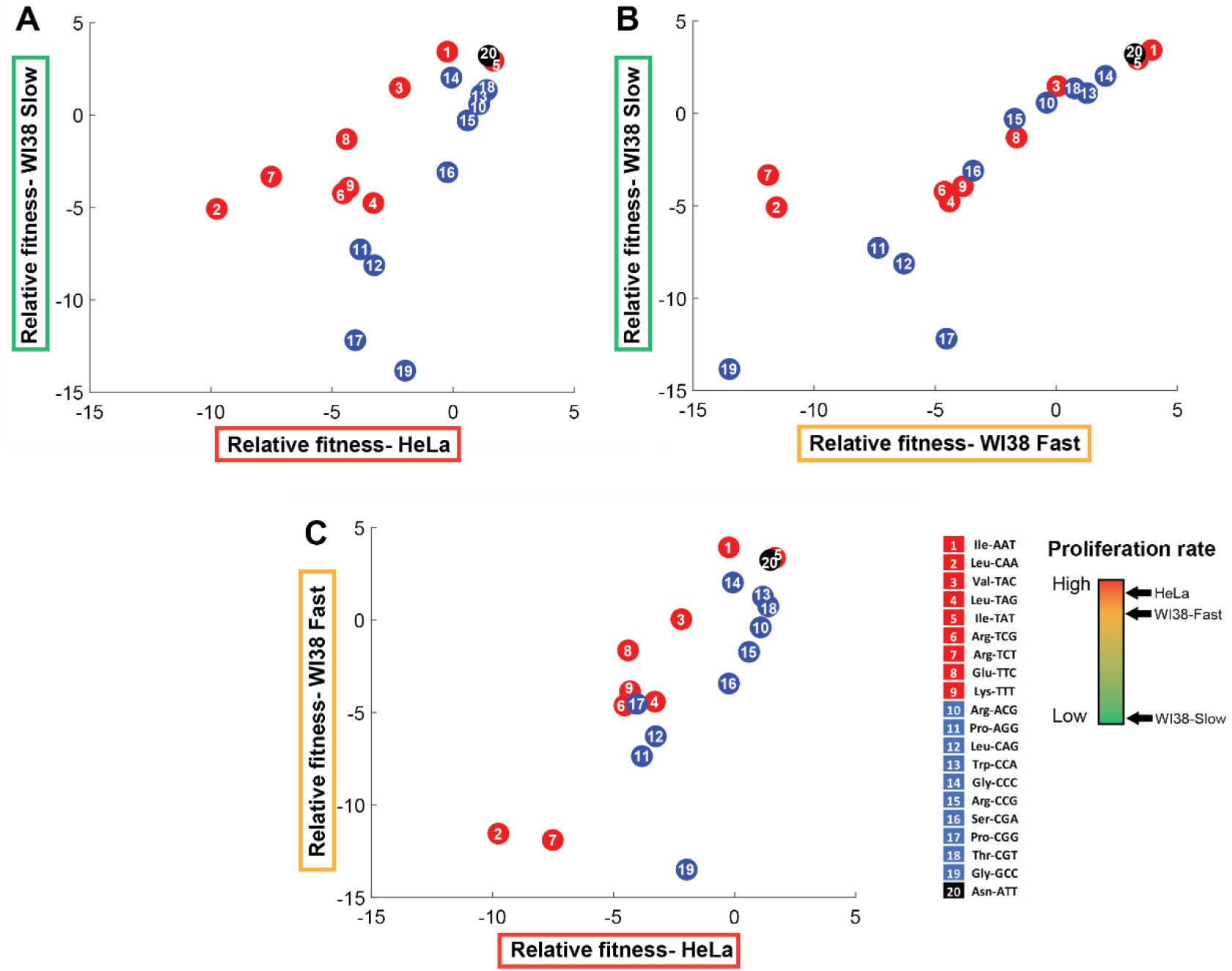
The tRNA essentiality depends on cell line origin and the proliferation rate. Scatter plots that compare the relative fitness of the edited tRNA cells between all pairwise combinations of three cell lines. The relative fitness of each CRISPR-targeted tRNA variant in each cell lines was determined based on day 7 of the cell competition, and was averaged over two biological repeats.

### CRISPR-targeting of tRNAs affects the transition from proliferative to arrested state

Having established the differential roles of tRNA for cellular proliferation, we turned to examine their essentiality in response to cell cycle arresting conditions. We focused on two distinct cell cycle arrest states-quiescence and senescence, which are reversible and irreversible G0 states, respectively.

To assess the role of tRNAs in entering these processes, we expressed the tRNA-CRISPR library described in Figures 1 and 3A in WI38 fast iCas9 cells. Besides the CRISPR system, we expressed in those cells a mCherry gene downstream to an endogenous promoter of human p21 (p21p-mCherry), a known marker for arrested cells. The p21 protein is a cyclin-dependent kinase inhibitor that promotes the entrance to cell cycle arrest. P21 is a primary mediator of the p53 pathway in response to DNA damage, which results in the loss of proliferation potential and induction of senescence (Abbas & Dutta 2009; Rufini et al. 2013). In addition, studies have shown that high p21 expression is essential for the transition to the quiescence state (Perucca et al. 2009; Wesley Overton et al. 2014).

We stably introduced a p21p-mCherry construct into WI38 fast iCas9 cells, followed by the creation of a clonal population that originated from a single cell. Then, we transduced the sgRNA library into the clonal WI38 fast iCas9 and p21p-mCherry cells. After that, we applied antibiotics selection, followed by induction of the iCas9 for 3 days to allow editing of the tRNA genes (Fig 5A). Then, we split the transduced population into three populations that were each allowed to grow continuously, yet under different conditions (see Materials and Methods). Quiescence was induced by growing the cells in a serum-free medium (Fig 5A, “Q” population). Senescence was induced by Etoposide at a sub-lethal concentration (Fig 5A, “S” population). The untreated population continued to grow in normal conditions (Fig 5A, “U” population). After two days, we measured the mCherry levels (normalized to the cell size, using the forward scatter (FSC) measure) of each of the three populations using a flow cytometer. We note that each population consists of 20-sub populations, one for each of the CRISPR-targeted tRNAs. We observed a significant difference in the mCherry/FSC distributions between the three populations, indicating differences in the regulation of p21 expression in response to the different types of arrest modes. The senescent population showed the widest distribution, with the highest mCherry/FSC value (Fig 5A). As expected, the untreated population showed the tightest distribution, with the lowest mCherry/FSC value (Fig 5A). The histogram of the quiescence population is similar to that of the untreated, with a small shift towards the higher mCherry/FSC values (Fig 5A).

**Figure 5.**
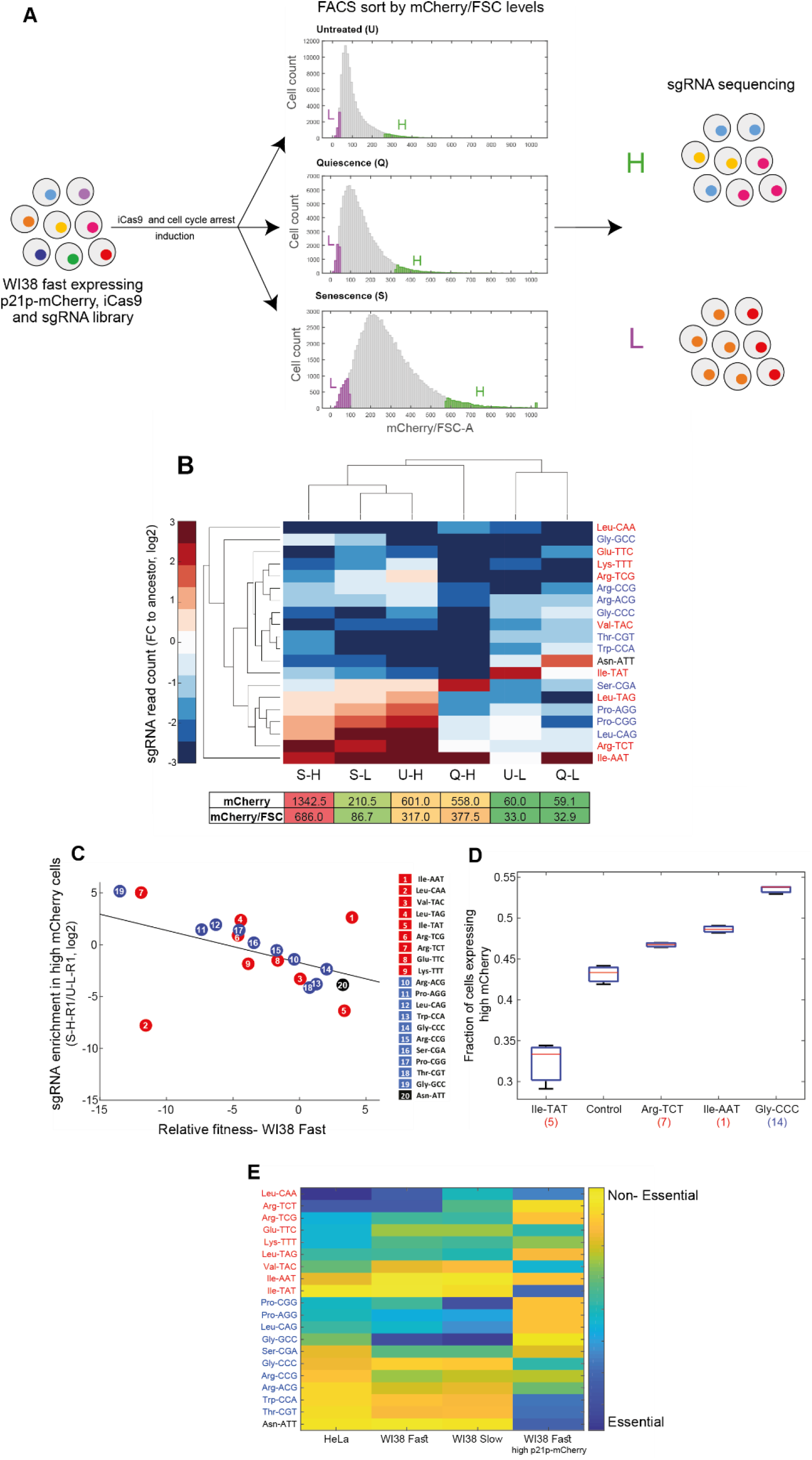
Essentiality of tRNAs for the response to cell cycle arresting signals. A| A procedure for multiplexed assay for tRNA essentiality for growth arrest. The tRNA-sgRNA library was transduced into WI38 fast cells that express iCas9 and mCherry gene under the endogenous promoter of human p21. After the induction of iCas9, we split the cell population (ancestor sample) into three populations, each treated with different conditions to stimulate the entrance to the cell cycle arrest state. After two days, we measured the mCherry/FSC levels of each population using FACS. The distributions of the three populations, based on the mCherry/FSC are shown in the middle (U-untreated; Q-quiescence; S-senescence). Then, each population was sorted according to the mCherry/FSC ratio, while separating the top and bottom 5% of each population (High bin-labeled in green; Low bin-labeled in purple). Then, we extracted the genomic DNA of each sample and deep sequenced the sgRNAs, looking for enriched and depleted sgRNAs (relative to the ancestor sample). B| Hierarchical clustering of the sorted samples and the sgRNAs based on the averaged changes (two biological repeats) in the sgRNA read count of each sorted sample normalized to the ancestor sample (log2). The lower table depicts the mean mCherry level and the mean mCherry/FSC ratio of each sorted population, based on the FACS measurements. C| A scatter plot comparing the essentiality of tRNAs to cellular proliferation and cell arrest in WI38 fast cells. Colors of tRNA families are as in previous figures. D| A box plot presenting the fraction of cells expressing high mCherry levels (>1700) in each CRISPR-targeted tRNA variant (three biological repeats). The control sample is a random sgRNA with no target sequence in the human genome. The number in parentheses on the x-axis denotes the gene copy number of each tRNA family. E | A heat map summarizing the essentiality of each tRNA to the different cell lines and proliferation states. Each row represents a tRNA family, classified to proliferation (red), differentiation (blue), or pseudo (black) tRNAs. Each column represents a cell line or condition. The color code depicts the essentiality of the tRNA. The essentiality is the z-transformed values of the (log2) Fold Change in sgRNA read count in each experiment (as described in Fig 3 and Fig 5C).

We could then progress towards a multiplexed assay for the essentiality of each of the 20 CRISPR-targeted tRNAs for responding to the signal and carrying out a program for cell arrest. We reasoned that cells targeted for tRNAs that are important for the transition from the base condition to a non-dividing state are likely to be underrepresented in the population of responding cells, i.e. cells with high p21p-mCherry levels. We have thus sorted each of the three populations based on the mCherry/FSC values, and sampled cells from the top and bottom 5% of each of the respective populations (i.e., High (‘H’) and Low (‘L’) bins, respectively). We hypothesized that the high mCherry/FSC samples will be enriched with responding cells, while the low mCherry/FSC samples will be enriched with non-responding cells (Fig 5A). From each sorted population we extracted the genomic DNA and deep-sequenced the sgRNAs (Fig 5A). We examined the diversity of the sgRNA read count in each sorted population of the different treatments, normalized to the sgRNA read count of the ancestor sample (the edited tRNA cell population, without any treatment). We thus obtained the normalized essentiality profile of 20 tRNAs at six samples (three conditions, times high and low mChery level for each). To assess the similarity of tRNA essentiality profiles of the six samples we used two-way hierarchical clustering of the samples and the tRNAs (Fig. 5B). Clustering samples across the 20 CRISPR-targeted tRNAs shows two main clusters. One cluster consists of all high mCherry/FSC samples from all three treatments (called, U-H, Q-H and S-H), as well as senescence low mCherry/FSC sample (i.e., S-L), suggesting that in the senescence treatment, even the Low mCherry/FSC cells, S-L sample, contains responding cells. The second cluster contains the two low mCherry/FSC samples from the untreated and quiescence populations (i.e., U-L and Q-L). Upon clustering the 20 CRISPR-targeted tRNAs across the samples we could detect some tRNAs that appear essential for entry into the arrested state. Interestingly, these tRNAs do not belong consistently to either the proliferation-tRNA or differentiation-tRNA sets. Yet, overall, same tRNAs tend to be more essential for both entries to quiescence and senescence.

We next compared the essentiality of the various tRNAs for the proliferation of WI38 fast cells (Fig. 3B) and their essentiality for entry into growth arrest and found a negative correlation throughout all but 2 of the tRNAs (Fig 5C, Pearson correlation, r = −0.45, p < 0.05). It seems that tRNAs that are essential for proliferation are relatively less essential for growth arrest. Thus, although the transition of cells into an arrested state has a more complex dependency on various sub-set of the tRNA pool it appears to necessitate tRNAs that are distinct from those needed for proliferation.

CRISPR-targeting of certain tRNAs triggers by itself, cell cycle arrest, independently of external induction. Under this alternative model, those tRNAs that appear to be most essential for growth arrest in response to the external signals actually trigger arrest to the lowest level upon their own targeting in otherwise-untreated cells. To address this concern, we examined the distribution of mCherry levels of certain CRISPR-targeted tRNA variants at the absence of any external arresting condition. We found that certain CRISPR-targeted tRNA variants (Arg-TCT, Ile-AAT and Gly-CCC) show elevated cell cycle arrest levels, as deduced from a higher fraction of high mCherry levels in those mutants compared to a control sample (which carries a random sgRNA without any target site in the human genome) (Fig 5D. CRISPR-targeted tRNA variants vs control sample, t-test, p < 0.05). On the other hand, genomic editing of Ile-TAT tRNA genes restricts the transition process, as inferred from the smaller fraction of high mCherry cells compared to the control sample (Fig 5D. targeted Ile-TAT sample vs control sample, t-test, p < 0.01). These results indicate that the editing of certain tRNAs indeed triggers, directly or indirectly, cell cycle arrest. In contrast, CRISPR-targeting of other tRNAs is crucial for the transition process from proliferative to arrested state, while growth itself is not dependent on these tRNAs.

Fig 5E shows a summary of tRNA essentiality across cell types and proliferation states. In general, we observed that while most of the proliferation tRNAs are essential in proliferating cell lines (HeLa, WI38-Fast and to a lesser extent in WI38-Slow), only part of the differentiation tRNAs is essential for these cell lines. Yet, we identified tRNAs whose targeting affected differently the different proliferative cell lines. For example, ArgTCT is highly essential to HeLa and WI38-Fast, while WI38-Slow cells are less dependent on this tRNA family. GlyGCC is mildly important for the growth of HeLa cells, while WI38-Fast and Slow cells are highly dependent on this tRNA family. Overall, the tRNAs can be classified into 4 main groups: tRNAs that are essential to all cell lines and proliferation states, such as LeuCAA; tRNAs that are essential to proliferative cell lines and less essential to the transition to arrested state, such as ArgTCG and ProAGG; tRNAs that are essential to transition from proliferation to arrest state, while proliferating cells are less dependent on them, as IleTAT and ThrCGT; tRNAs that are relatively dispensable in all cell lines and proliferation states, like IleAAT. Interestingly, the pseudogene tRNA AsnATT appears dispensable in all proliferative assays but essential for the translation into cell cycle arrest.

## Discussion

In this work, we aimed to decipher the causal relation between the tRNA expression and the cellular state in human cells. For that, we manipulated the cellular tRNA pool in various cell lines and proliferation states by targeting 20 different tRNA familes using CRISPR-iCas9. We found that cells from different origins and proliferation rates are dependent on distinct sets of tRNAs. Mainly, we identified that proliferation tRNAs, as those whose CRISPR-edited are deleterious to highly proliferating cells. We found that slowly proliferating fibroblasts are more sensitive to the previously defined ‘differentiation tRNAs’ (Gingold et al. 2014). However, here we found certain tRNAs that upon CRISPR-editing do not behave as expected by their original proliferation-differentiation classification. In particular tRNAs IleAAT, IleTAT, and ValTAC, which were originally defined as proliferation tRNAs are no longer assigned to that category, while ProCGG, ProAGG, and LeuCAG that were originally regarded as differentiation tRNAs, should now be considered as proliferation tRNAs since they are essential for cellular proliferation. Additionally, we explored how the expression manipulation of tRNA families affects the ability of proliferative cells to respond to cell cycle arresting signals by entering into quiescence or senescence. We found that several tRNAs that are essential for growth are relatively dispensable for the entry into cell arrest, while tRNAs that are less essential for growth were more likely to be needed for mediating the arresting response to external stimuli. Indeed, we observed a negative correlation, across the 20 manipulated tRNAs, between their essentiality in growth and in cell arrest (Fig 5C). Yet, the actual identity of the tRNAs that are essential for cell arrest does not map precisely onto the distinction between the two tRNA sets. We note, though, that the previously defined “proliferation tRNAs” probably included tRNAs that serve in the translation of genes that carry out functions broader than mere cell division, e.g. transcription, translation, etc. These cellular functions are surely needed in additional cellular states such as when cells enter an arresting state (Hernandez-Segura et al. 2017; Casella et al. 2019).

The set of proliferation tRNAs also serves the translation of proteins that are not necessarily involved only in cellular proliferation, but more generally with cell-autonomous functions, such as gene expression. It is thus natural that the process of cell cycle arrest might also depend on some of these tRNAs that contribute to the translation of genes that carry out such cell-autonomous functionalities.

Changes in the expression of tRNAs in cancer are well established. Furthermore, there are several lines of evidence that point to causal effects of tRNAs in cancerous growth and metastasis formation (Felton-Edkins et al. 2003; Pavon-Eternod et al. 2009; Pavon-Eternod et al. 2013; Goodarzi et al. 2016). Indeed, several studies showed the potential of tRNAs as cancer biomarkers (Zhang et al. 2018; Hernandez-Alias et al. 2020). Our current results naturally raise the possibility that tRNAs can become also new targets for therapeutic strategies. Downregulation of certain tRNAs may serve as a growth arresting treatment in the context of cancer. CRISPR-editing of specific tRNAs was found here to have a close-to-immediate halt of proliferation, even in the extreme case of HeLa cells. Crucially, our results show that some tRNAs are essential specifically for cancerous cells and not in differentiated cells. Such selective essentiality is important as it may suggest that targeting these tRNAs in a mixture of healthy and cancerous cells may affect mainly cancer. It is worth mentioning in that respect the continuous improvement in precise delivery and activation of CRISPR technology in-vivo (Shim et al. 2017; Tong et al. 2019), such development increase the prospects of manipulation of specific tRNAs in a target within the body.

## Materials and methods

### SgRNA design and cloning

sgRNA candidates targeting various tRNA families were designed by providing tRNA sequence to “http://chopchop.cbu.uib.no/” (Montague et al. 2014). For each tRNA family, analyses were done for each of its unique sequences in the human genome. Only sgRNA candidates targeting the anticodon loop were considered. Then, a single sgRNA was chosen for each tRNA family that was predicted to target the maximum number of genomic copies.

“Restriction-free” cloning was performed to create sgRNA plasmids for targeting tRNA families. For each sgRNA sequence, long primers were ordered and used as megaprimers (sup data, table1). The PCR reaction was conducted using iProof master mix (X2) (Bio-Rad; 172-5310), 1-1.5μl of Forward and Reverse primers and 50ng of lenti-sgRNA plasmid (Addgene; 52963) in a 50μl total volume reaction, for 12 cycles, annealing: Tm = 60°C for 20sec, elongation: 72°C for 10 min. To eliminate the original plasmid, PCR products were incubated with 1μl DpnI enzyme (NEB; R0176S) for 1 hour at 37°C, and then 20 min at 80°C for inactivation. Following the DpnI treatment, plasmids were transformed into DH5α competent bacteria using standard heat shock transformation technique (Sambrook & Russell 2001). To find recombinant plasmid, colonies that grow under ampicillin selection were tested by sequencing of the purified plasmid using Wizard Plus SV Minipreps DNA Purification (Promega; A1330). Primer used for the validation sequencing is TTAGGCAGGGATATTCACCA. For massive plasmid purification, NucleoBond Xtra Midi kit (Macherey-Nagel; 740412.50) was used.

### Cell culture

293T cells (ATCC; CRL-3216) were grown in DMEM high glucose (Biological Industries; 01-052-1A) supplemented with 10% FBS, 1% penicillin/ streptomycin (P/S) and 1% L-Glutamine.

HeLa cells (kindly given by Prof. George Church’s lab) were grown in DMEM + NEAA (Life Technologies; 10938-025) – Dulbecco’s Modified Eagle Medium with 4.5mg/ml D-Glucose and Non-Essential amino acids. Supplemented with 10% FBS, 1% penicillin/ streptomycin (P/S), 1% L-Glutamine and 1% Sodium Pyruvate.

WI38 Slow and fast fibroblasts (referred to as WI-38/hTERT^slow^, 48 PDL and WI-38/hTERT^fast^, 484 PDL (Milyavsky et al. 2003) were grown in MEM-EAGLE + NEAA– Earle’s salts base with Non-Essential amino acids (Biological Industries; 01-025-1A), supplemented with 10% FBS, 1% penicillin/ streptomycin (P/S) and 1% L-Glutamine.

For iCas9 plasmid selection, 200μg/ml Hygromycin (Thermo Fisher; 10687010) were added to the medium and refreshed every two days. For sgRNA plasmid selection, 2μg/ml Puromycin (Sigma-Aldrich; 58-58-2) were added to the medium of HeLa (Fig 2–4). For sgRNA plasmid selection in fitness assay performed on WI38 slow and fast cell lines (Fig 4), 2μg/ml Puromycin were added to the medium, although these cell lines had already puromycin resistance from their immortalization process. 10 μg/ml Blasticidin (InvivoGen; BLL-38-02A) were added to the medium of WI38 fast for sgRNA plasmid selection in cell cycle arrest treatment assay (Fig 5). For stable transduction, the medium containing antibiotics was refreshed every two days, for all cell lines. For iCas9 induction, 1μg/ml Doxycycline (Sigma-Aldrich; 10592-13-9) was added to the medium and refreshed every two days.

### Generation of stable cell-line

#### Generation of iCas9 carrying cell lines

HeLa cells carrying iCas9 were generously provided by Prof. George Church.

WI38 slow and WI38 fast were seeded onto 10cm plates such that cell confluence will be approximately 70% the next day. 5μg of iCas9 vector (pB-Cas9 & pB-support vector) was transfected to cells with fresh MEM-EAGLE medium (5ml) and 15μl of Poly-jet transfection reagent (SignaGen; SL100688). After four hours, the medium was replaced with a fresh MEMEAGLE medium. Five days after the transfection, MEM medium containing 200 μg/ml Hygromycin was added to the transfected cells, refreshed every day for approximately one month.

#### Generation of iCas9 cells with sgRNA plasmids

##### Step 1 – viral vector production

A 10 cm plates were covered with ~2ml Poly-L-Lysine (Sigma; P4707) and then 293T cells (ATCC; CRL-3216) were seeded onto the covered 10 cm plates such that cell confluence will be approximately 70% the next day. A day after, 2.5μg of PMD2.G (Addgene;12259) and 10.3μg psPAX2 (addgene;12260) packaging vectors were co-transfected with 7.7μg of the appropriate sgRNA plasmids using 40μl of jetPEI (Polyplus; 101-10N) in DMEM high glucose medium (5ml). Note that for the pooled sgRNA experiments, the sgRNA plasmids were added in equal amounts. After 48 and 72 hours, virus-containing medium was collected and centrifuged for 15 minutes at 3200g, 4°C. Supernatant was collected to new tube, and 1.25ml PEG solution from PEG virus precipitation kit (BioVision; K904-50/200) was added. The virus containing tube was stored in 4°C for at least 12 hours (over-night). Virus-contained tubes were centrifuge for 30 min at 3200g, 4°C. Supernatant was removed and the virus pellet was suspended with 100μl virus resuspension solution from PEG virus precipitation kit.

##### Step 2 – cell transduction

WI38 Slow, WI38 Fast and HeLa cells were seeded onto 10 cm plates cells such that cell confluence will be approximately 50% the next day. On the day of transduction (24 hours after cell seeding), cell’s medium was replaced with 5μg/ml Poly-Brene (Sigma; TR-1003) contained medium (5ml). Suspended viruses were added to each plate according to the calibrated titer load (MOI ~ 0.3).

#### Generation of WI38 fast iCas9 cell lines carrying p21p-mCherry

The fragment containing human p21 promoter-mCherry-3NLS from the plasmid bank of the Weizmann Institute (originally created by Prof. Moshe Oren’s lab) replaced the U6 region of lenti-sgRNA plasmid (Addgene; 52963) using Gibson Assembly cloning (Gibson Assembly Master Mix, NEB; E2611). After plasmid extraction, viruses containing the plasmid were produced and WI38 fast-iCas9 cells were transduced with the viruses as described above. After a week of antibiotics selection, the cells were sorted using Flow cytometer into single cells and seeded in 96 well plate containing condition media. Flow cytometry analysis was performed on a BD FACSAria Fusion instrument (BD Immunocytometry Systems) equipped with 488-, 405-, 561- and 640-nm lasers, using a 100-μm nozzle, controlled by BD FACS Diva software v8.0.1 (BD Biosciences). Further analysis was performed using FlowJo software v10.2 (Tree Star). mCherry was detected by excitation at 561 nm and collection of emission using 600 LP and 610/20 BP filters. The described experiments were done in one of the single clones of WI38 fast iCas9+ p21p-mCherry.

### Evaluating the tRNA expression levels and the editing gene variants of CRISPR-targeted tRNAs (Fig 2)

#### Experimental procedure

48 hours after lentiviral infection, containing a single sgRNA plasmids, the cell’s media was replaced with antibiotics-containing medium (2μg/ml Puromycin), to allow selection of infected cells, for 7 days. Then, cells were grown for 12 days in medium contained both antibiotics and 1μg/ml doxycycline, for selection and iCas9 expression respectively. During the time course, the cells were diluted every 2 days in a ratio of 1:2.5. A cell sample was taken every 3 days and frozen at −80°C. in addition, HeLa iCas9 cells without sgRNA (referred as WT cells) were grown with doxycycline and sampled as described above.

#### Mature tRNA sequencing-library preparation and data processing

tRNA sequencing protocol was adapted from (Zheng et al. 2015) with minor modification. Small and large RNA were extracted from frozen samples using NucleoSpin miRNA kit (Macherey-Nagel; 740971.50). 1 ug RNA was mixed with 0.025 pmole of tRNA standards (e.coli-tyr-tRNA and s.cerevisia-phe-tRNA at ration of 1:8 (Sigma-Aldrich; XX)). Reverse transcription was done using TGIRT™-III Enzyme (InGex; 5073018), with the indicated primers. 3’ adaptor was ligated to the cDNA using T4 ligase (NEB; M0202S). The cDNA was purified using Dynabeads myOne SILANE (life Technologies; 37002D) after each step. The library was amplified using NEBNext High-Fidelity 2X PCR Master Mix (NEB; M0541S) and cleaned using SPRI-beads with left-side size selection protocol. Samples were pooled and sequenced using a 75bp single read output run on MiniSeq high output reagent kit (Ilumina; FC-420-1001).

Primer Reverse-transcription DNA 5’-CACGACGCTCTTCCGATCTT −3’ Reverse-transcription RNA 5’-rArGrArUrCrGrGrArArGrArGrCrGrUrCrGrUrG-3’ 3’-ligation adaptor 5’-AGATCGGAAGAGCACA-3’ Read were trimmed using homerTool. Alignment was done to the genome and mature tRNA using Bowtie2 with parameters --very-sensitive-local. Read aligned with equal alignment score to the genome and mature tRNA were annotated as mature tRNA. Reads aligned to multiple tRNA genes were randomly assigned when mapping to identical anticodon, and discarded from the analysis if aligned to different anticodon. Read count was done using BedTools-coverage count. Variant calling for detection of mutation and indels was done using samtools command “mpileup” with the parameters: “-A -q1 -d100000”.

The fold change in tRNA expression for each tRNA isoacceptor was calculated as follows: the normalized read counts for each tRNA isoacceptor were summed up over all tRNA genes in each sample (treated and WT cells) and in each time point along the iCas9 induction four time points. Then, a ratio of the summed read count between the treated and WT samples was calculated for each time point. This procedure was done for each of the two biological repeats. The fraction of edited tRNA reads was calculated as follow: for each tRNA isoacceptor, the number of aligned reads with mutations were summed up for all tRNA genes, and normalized to the sum of maximum coverage for all tRNA gene in the vicinity of the mutation. Then, a ratio of the fraction of edited reads between the treated and WT samples was calculated for each time point.

#### Genomic tRNA sequencing – library preparation and data processing

Genomic DNA was purified from frozen cell samples of the sgRNA competition experiment using PureLink genomic DNA mini kit and used as template for PCR to amplify specifically the tRNA isodecoder genes of the CRISPR-targeted tRNAs in the population. PCR reaction was conducted using KAPA HiFi HotStart ReadyMixPCR (X2) (kapabiosystems; KK2601), 10μM of each primer and 100ng of genomic DNA extracted from the samples in a 60μl total volume reaction, for 25 cycles. Primers and annealing temperatures are listed (sup data, table 2). After PCR validation using agarose gel, the PCR product was purified with SPRI-beads (Agencourt AMPure XP, Beckman Coulter; A63881), using left-side size selection protocol while the PCR product and beads were mixed at 1:1.5 ratio. Then, each sample was barcoded with a different Illumina barcode while the Forward primer was fixed and the reverse primer included the different barcodes. The barcoding PCR reaction that was conducted using KAPA HiFi HotStart ReadyMixPCR Kit (X2), 10μM of Illumina primers and 0.35-0.7ng of PCR product in a 10μl total volume reaction, for 15 cycles. Additional clean-up step was performed using SPRI beads with left-side size selection protocol while the PCR product and beads were mixed at 1:1.5 ratio. Samples were pooled in equal amounts and sequenced. We performed a 75bp single read output run on MiniSeq high output reagent kit (Ilumina; FC-420-1001).

After the reads were trimmed using Cutadapt, we used CRISPResso (Pinello et al. 2016) to quantify NHEJ events in the CRISPR-Cas0 targeted samples. For the pipeline analysis, we added the WT amplicon sequences as well as the sgRNA sequences. The minimum identity score for the alignment was set to 0. For each sample, we quantified the read count with WT amplicon or amplicons with NHEJ events. For the reads with NHEJ events, we quantified the reads with Insertions and deletions.

The fraction of edited tRNA reads was calculated as follow: for each tRNA isoacceptor, the number of edited reads were summed up for all tRNA genes, and normalized to the sum of aligned reads (edited and WT reads). Then, a ratio of the fraction of edited reads between the treated and WT samples was calculated for each time point.

### Evaluating the fitness of tRNA knockout variants – pooled sgRNAs (Fig 3–4)

#### Experimental procedure

48 hours after lentiviral infection, containing the pooled sgRNA plasmids, the cell’s media was replaced with antibiotics-containing medium, to allow selection of infected cells (2μg/ml Puromycin). After 24 hours of selection, cells were grown for 14 days in medium contained both antibiotics and 1μg/ml doxycycline, for selection and iCas9 expression respectively. During the time course, the cells were diluted every 2 days in a ratio of 1:2.5. A cell sample was taken every 3 or 4 days and frozen at −80°C.

#### sgRNA sequencing-library preparation and data processing

Genomic DNA was purified from frozen cells samples of the sgRNA competition experiment using PureLink genomic DNA mini kit (Invitrogene; K182000) and used as templates for PCR to amplify specifically the sgRNAs in the population. PCR reaction was conducted using iProof master mix (X2), 10μM of each primer and 20ng of genomic DNA extracted from the samples in a 50μl total volume reaction, for 26 cycles, Tm = 64°C. The primers used to amplify the sgRNA region were:

Forward primer-GCTTACCGTAACTTGAAAGTATTTCGATTTCTTGG

Reverse primer-CTTTTTCAAGTTGATAACGGACTAGCCTTATTTTAAC

After PCR clean-up using Wizard SV Gel and PCR Clean-Up (Promega; A9281), samples were run in 2% agarose gel to ensure that the PCR product is compose of a single amplicon in the appropriate size. Next, Hiseq libraries were prepared using the sequencing library module from (Blecher-Gonen et al. 2013). Briefly, blunt ends were repaired, Adenine bases were added to the 3’ end of the fragments, barcode adapters containing a T overhang were ligated, and finally the adapted fragments were amplified. The process was repeated for each sample with a different Illumina DNA barcode for multiplexing, and then all samples were pooled in equal amounts and sequenced. We performed a 125bp paired end high output run on HiSeq 2500 PE Cluster Kit v4. Base calling was performed by RTA v. 1.18.64, and de-multiplexing was carried out with Casava v. 1.8.2, outputting results in FASTQ format.

De-multiplexed data was received in the form of FASTQ files split into samples. First, SeqPrep (https://github.com/jstjohn/SeqPrep) was used to merge paired reads into a single contig, to increase sequence fidelity over regions of dual coverage. The size of each contig was then compared to the amplicon’s length. Next, the forward and reverse primers were found on each contig (allowing for 2 mismatches) and trimmed out. This step was performed for both the forward and reverse complement sequences of the contig, to account for non-directional ligation of the adaptors during library preparation. After the primers were trimmed, the contig was tested again for its length to ensure no Indels had occurred. Contigs were then compared sequentially to all sgRNAs, comparing the sequence of each contig to the sequence of each sgRNA. Any contig without a matching sgRNA within two mismatches or less was discarded. Contigs with more than a single matching sgRNA with the same reliability were also discarded due to ambiguity. Each contig that passed these filters was counted in a keyvalue data structure, storing all sgRNA types and their frequency in each sample.

The relative fitness of each CRISPR-targeted tRNA variant was estimated by calculating the fold change of the sgRNA frequency in each time point relative to day 0 (before adding Doxycycline). We chose to explore the relative fitness of the tRNA knockouts based on relative frequency of their targeting sgRNAs at day 7 (relative to day 0), due to the dynamics of the cell population composition along the iCas9 induction. In the early days of the induction, the iCas9 activity does not reach saturation (Yuen et al. 2017), thus the cell population is dominated by partially CRISPR-edited cells. In the late days of the iCas9 induction, the less fit CRISPR-targeted tRNA cells might be eliminated from the population due to a negative selection, a process that can result in lower frequency of the tRNA-edited cells. Nonetheless, the relative fitness estimated by the different days of the competition is highly correlated (Fig S3A).

### Evaluating the essentiality of edited-tRNA variants for entering into cell cycle arrest state – (Fig 5)

#### Experimental procedure

Clonal WI38 fast iCas9 + p21p-mCherry cells were transduced with the sgRNA library as described above. After 6 days of antibiotics selection (10 μg/ml Blasticidin), the iCas9 was induced using 1μg/ml doxycycline for 3 days (ancestor sample). Then, the cells were split into three populations. One population continued to grow in normal conditions (untreated sample). The second population was grown with serum-free media-0% FCS (quiescence sample). The third population was grown with media containing 2.5uM Etoposide (Sigma-Aldrich; 33419-42-0) (senescence sample). After two days with treatment, the cells were sorted using Flow cytometer according to the mCherry/FSC levels, while sorting the top and bottom 5% of the population. FACS paramters as described above (see “stable cell line generation”)

#### sgRNA sequencing-library preparation and data processing

From each sorted population, together with the ancestor sample, the genomic DNA was extract using PureLink genomic DNA mini kit and used as templates for PCR to amplify specifically the sgRNAs in the population. PCR reaction was conducted using 2X KAPA HiFi HotStart ReadyMix, 10μM of each primer and 20ng of genomic DNA extracted from the samples in a 50μl total volume reaction, for 20 cycles, Tm = 58°C. We used shifted primers to increase library complexity (sup data, table 1). The PCR products were purified with SPRI-beads using left-side size selection protocol while the PCR product and beads were mixed at 1:1.5 ratio. The barcoding PCR and final PCR clean-up was done as described for the genomic tRNA library preparation. Samples were pooled and sequenced using a 75bp single read output run on MiniSeq high output reagent kit (Ilumina; FC-420-1001).

Reads were trimmed using cutadapt and then clustered into unique sequences using vsearch. Each unique read was then aligned to the matched sgRNA sequence, allowing up to two mismatches. Finally, we stored all sgRNA types and their frequency in each sample.

### Evaluating the essentiality of edited tRNA variants for entering into cell cycle arrest state – single sgRNAs (Fig 5D)

Clonal WI38 fast iCas9 + p21p-mCherry cells were transduced with single sgRNAs as described above. After 6 days of antibiotics selection (10 μg/ml Blasticidin), the iCas9 was induced using 1μg/ml doxycycline for 3 days. Then, the mCherry levels of each cell line were analyzed using Attune Flow Cytometer.

## Acknowledgments

We wish to thank Tammy Biniashvili and Omer Asraf for the help with the deep-sequencing analyses. We thank Dr. Tomer Meir Salame from The Weizmann Institute of Science’s Flow Cytometry Core Facility for the help with the Flow Cytometry experiments. We thank Lior Roitman and Dr. Hilah Gal from Valery Krizhanovsky group from the Weizmann Institute for the help and guidance with the cell cycle arrest experiments. A special thank for Dr. Hila Gingold and the Pilpel lab, for the stimulating discussions.

## Supplementary figures

**Fig S1.**
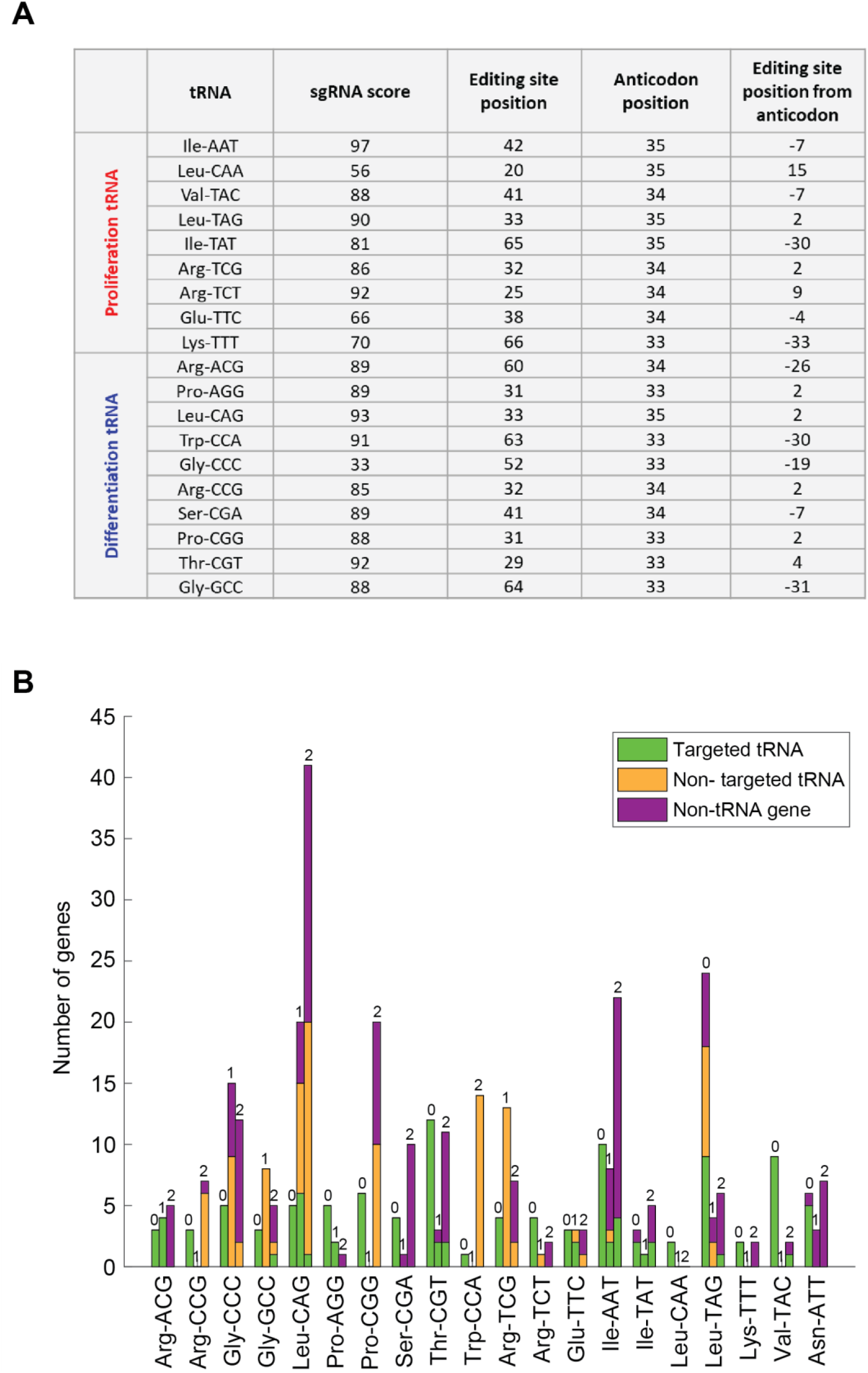
sgRNA targeting parameters and off-targets potential. A| A table representing various parameters of the sgRNA library. The sgRNA parameters include the score for each sgRNA as well as the editing position. The sgRNA score was determined using CHOPCHOP, a CRISPR web toolbox (Montague et al. 2014). The editing position is illustrated using the editing site position (relative to the start of the tRNA gene), the anticodon position (the first nucleotide of the anticodon relative to the tRNA gene start), and the editing site position relative to the anticodon. B| A bar plot representing the targets of the sgRNA library. Each three grouped bars depict potential on-targets and off-targeted with different degrees of sequence similarities between their sequence and the sgRNAs. The left bar in each triplet of grouped bars depicts the number of targets with full complementarity to the sgRNA sequence (marked as ‘0’ mismatches). The middle bar depicts the number of targets and off-targets with one mismatch (marked with ‘1’) and the right bar depicts the number of targets and off-targets with two mismatches (marked with 2). The colors depict the identity of the targeted gene: Green- a tRNA gene that belongs to the CRISPR-targeted tRNA family. Orange-a tRNA gene that belongs to unCRISPR-targeted tRNA family. Purple-a non-tRNA gene. The last two gene types are considered as off-targets

**Fig S2.**
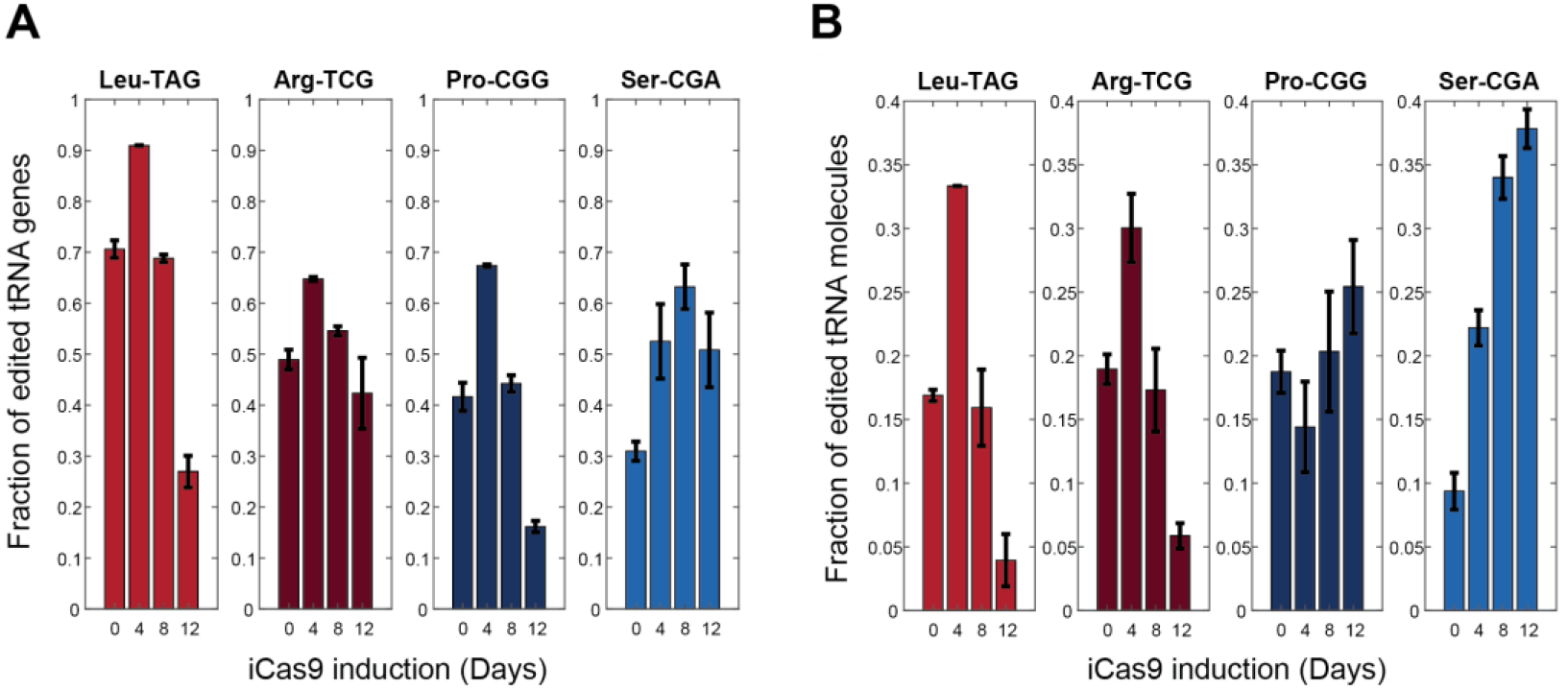
Fraction of edited tRNAs in CRISPR-targeted HeLa cells. Bar plots describing the genomic and mature tRNA sequencing results of CRISPR-targeted tRNA HeLa cells. A| Genomic tRNA sequencing: The fraction of the edited tRNA genes from the total tRNA reads (see Materials and Methods). B| Mature tRNA sequencing: The fraction of the edited tRNA molecules from the total tRNA reads (see Materials and Methods). Each bar represents a time point during the iCas9 induction. The two left plots describe the proliferation CRISPR-targeted tRNA samples and the two left plots describe the differentiation CRISPR-targeted tRNA samples.

**Fig S3.**
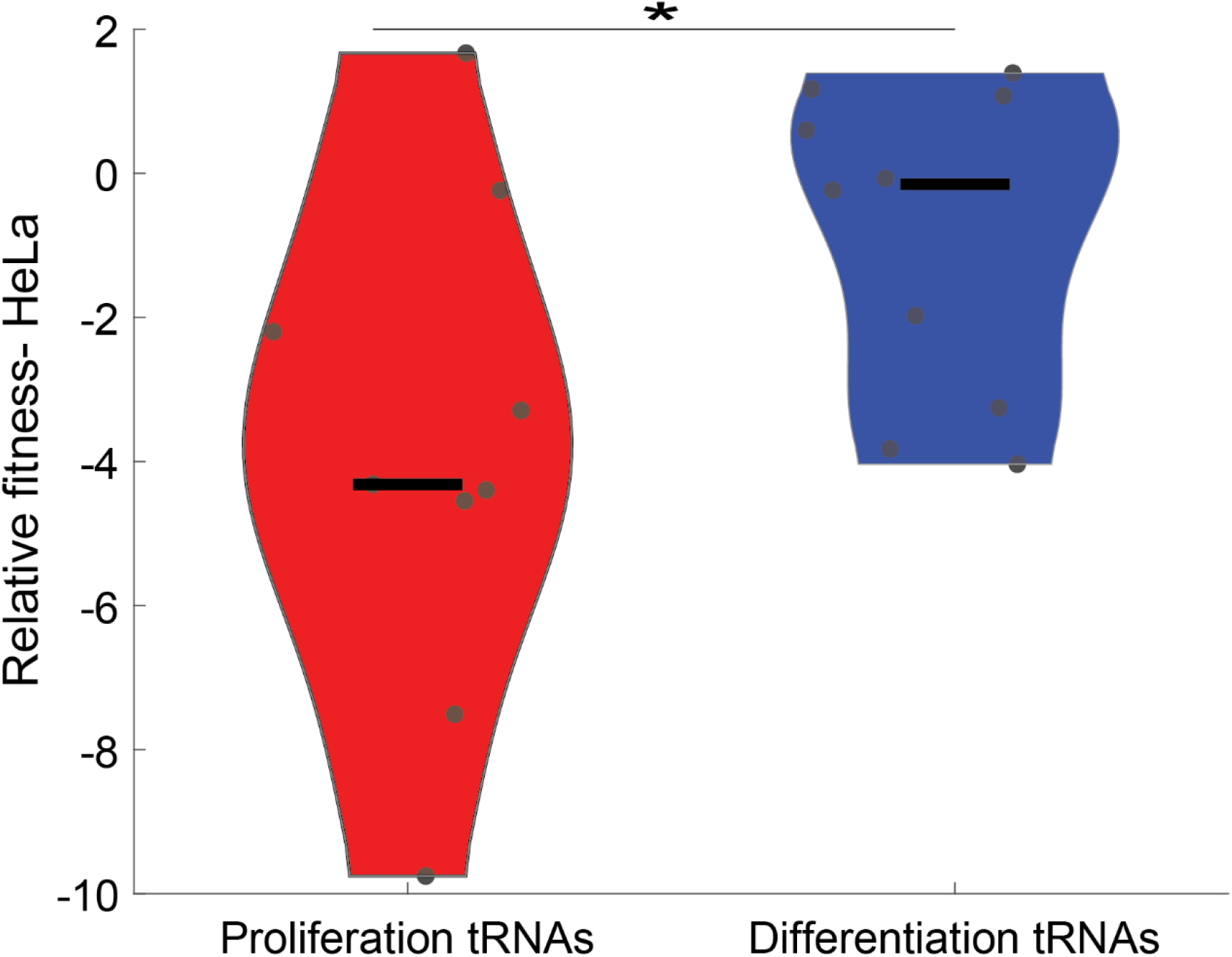
Relative fitness of targeted tRNA variants in HeLa cells. A violin plot describing the difference in the relative fitness of targeted tRNA variants in HeLa cells between proliferation and differentiation tRNAs (* – Wilcoxon rank-sum test, p < 0.05). The relative fitness on the y-axis is calculated as described in Figure 2. Each dot depicts a single tRNA family.

**Fig S4.**
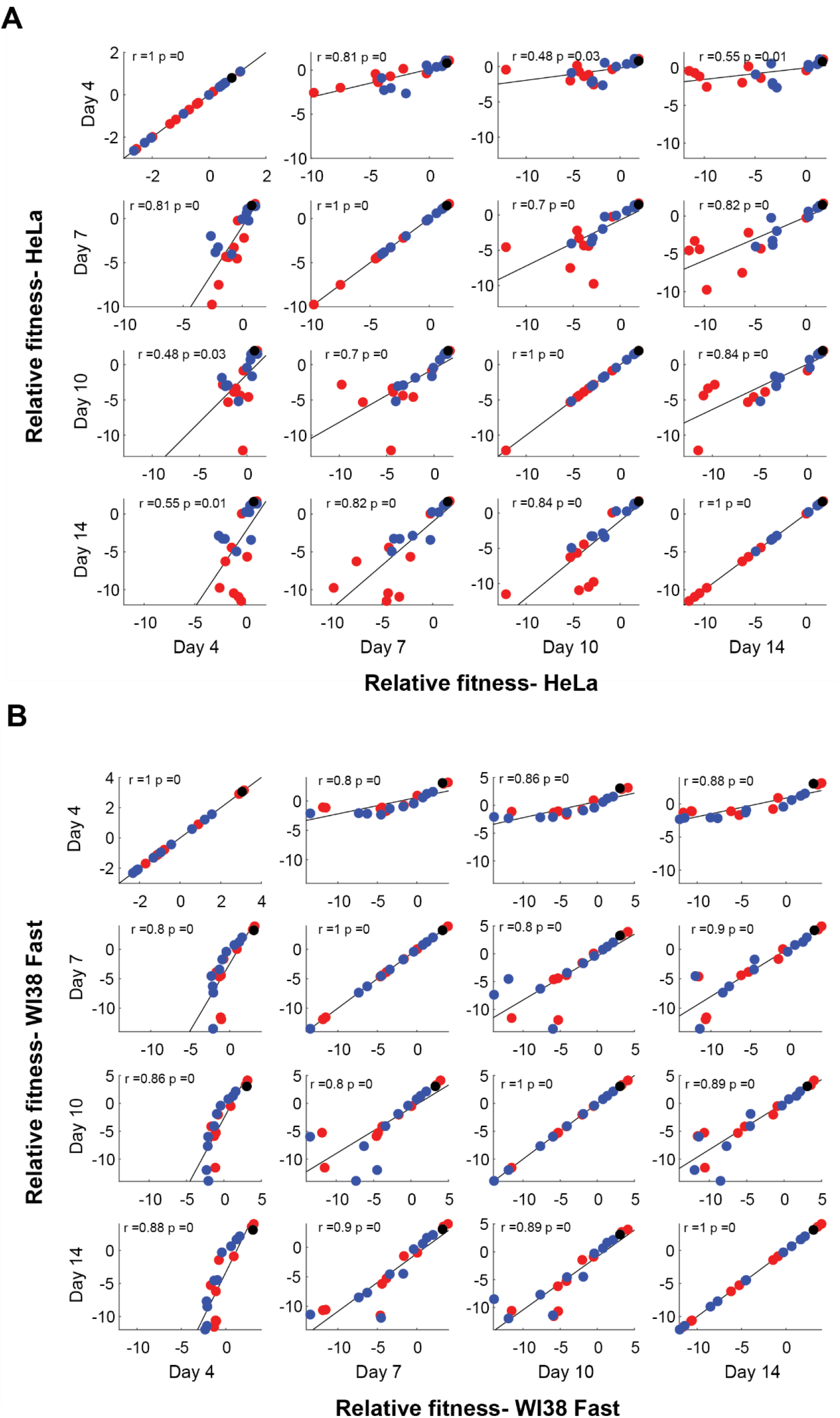
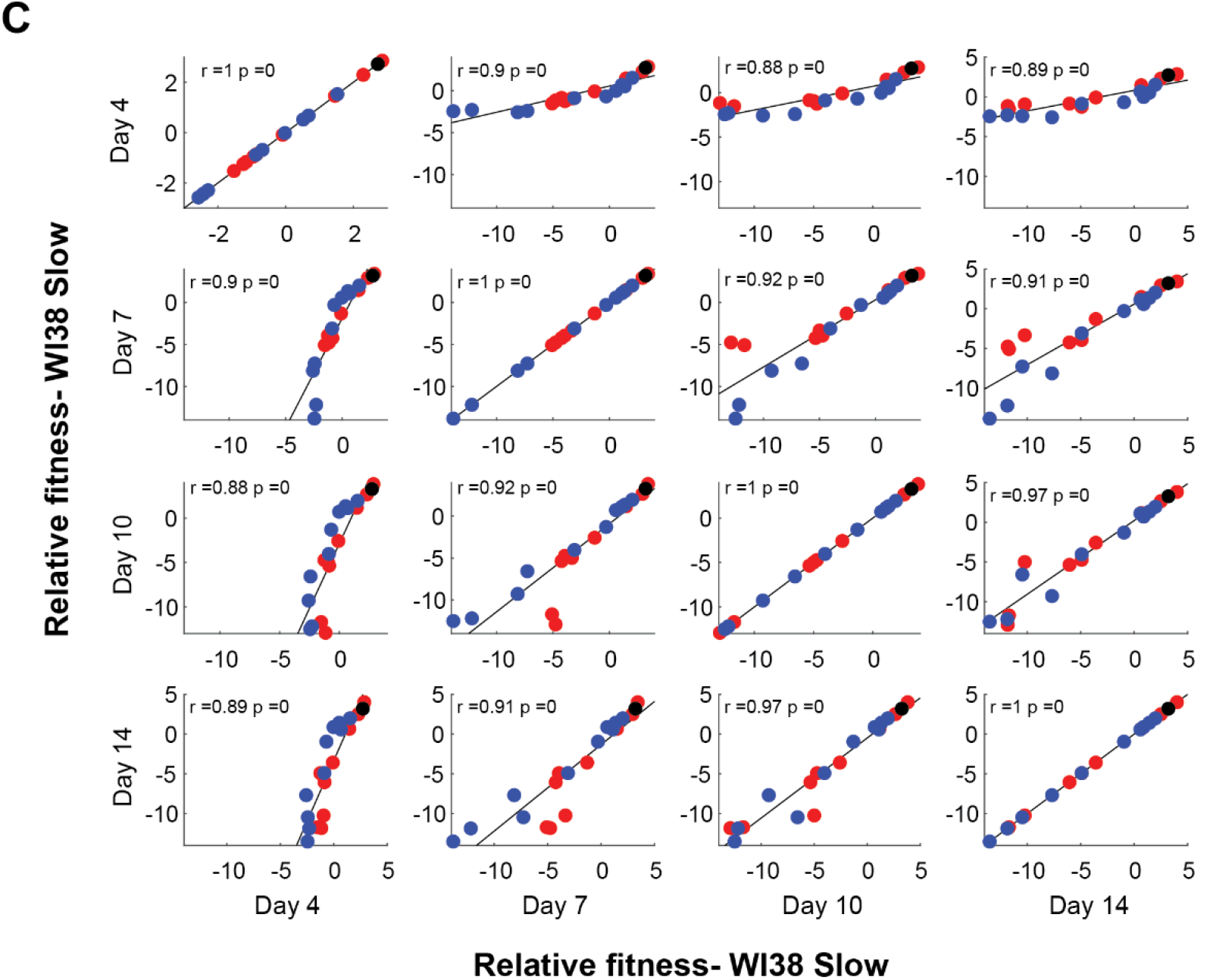
The correlative relationships between the relative fitness of CRISPR-targeted tRNA variants at different time points along the cell competition. Scatter plots representing the correlation between the relative fitness of the CRISPR-targeted tRNA variants in different days along the cell competition (Day 4, 7, 10, and 14). The colors depict the tRNA family (Red-proliferation tRNAs; blue-differentiation tRNAs; black-pseudo tRNA). The correlation coefficient r and p-values are detailed for each scatter plot. A| HeLa cells B| WI38 Fast cells C| WI38 Slow cells.

## References

Abbas, T. & Dutta, A., 2009. P21 in cancer: Intricate networks and multiple activities. Nature Reviews Cancer, 9(6), pp.400–414.

Aviner, R. et al., 2015. Uncovering Hidden Layers of Cell Cycle Regulation through Integrative Multi-omic Analysis. PLoS Genet, 11(10), p.1005554. Available at: https://journals.plos.org/plosgenetics/article/file?id=10.1371/journal.pgen.1005554&type=printable [Accessed June 18, 2019].

Bar-joseph, Z. et al., 2008. Genome-wide transcriptional analysis of the human cell cycle identifies genes differentially regulated in normal and cancer cells., 105(3).

Besson, A., Dowdy, S.F. & Roberts, J.M., 2008. CDK Inhibitors: Cell Cycle Regulators and Beyond. Developmental Cell, 14(2), pp.159–169.

Blecher-Gonen, R. et al., 2013. High-throughput chromatin immunoprecipitation for genomewide mapping of in vivo protein-DNA interactions and epigenomic states. Nature protocols, 8(3), pp.539–54.

Bludau, I. & Aebersold, R., 2020. Proteomic and interactomic insights into the molecular basis of cell functional diversity. Nature Reviews Molecular Cell Biology, pp.1–14. Available at: http://www.nature.com/articles/s41580-020-0231-2.

Casella, G. et al., 2019. Transcriptome signature of cellular senescence. Nucleic acids research, 47(14), pp.7294–7305.

Cheung, T.H. & Rando, T.A., 2013. Molecular regulation of stem cell quiescence. Nature Reviews Molecular Cell Biology, 14(6), pp.329–340.

Collado, M. & Serrano, M., 2010. Senescence in tumours: Evidence from mice and humans. Nature Reviews Cancer, 10(1), pp.51–57.

Dittmar, K.A., Goodenbour, J.M. & Pan, T., 2006. Tissue-Specific Differences in Human Transfer RNA Expression. PLoS Genetics, 2(12). Available at: http://journals.plos.org/plosgenetics/article/file?id=10.1371/journal.pgen.0020221&type=printable [Accessed January 10, 2018].

Felton-Edkins, Z.A. et al., 2003. The mitogen-activated protein (MAP) kinase ERK induces tRNA synthesis by phosphorylating TFIIIB, Available at: https://www.embopress.org/doi/pdf/10.1093/emboj/cdg240 [Accessed June 26, 2019].

Frumkin, I. et al., 2018. Codon usage of highly expressed genes affects proteome-wide translation efficiency. Proceedings of the National Academy of Sciences of the United States of America, 115(21), pp.E4940–E4949. Available at: www.pnas.org/lookup/suppl/doi:10.1073/pnas.1719375115/-/DCSupplemental.www.pnas.org/cgi/doi/10.1073/pnas.1719375115 [Accessed April 3, 2020].

Gardin, J. et al., 2014. Measurement of average decoding rates of the 61 sense codons in vivo. eLife, 3, pp.1–20.

Gingold, H. et al., 2014. A Dual Program for Translation Regulation in Cellular Proliferation and Differentiation. Cell, 158(6), pp.1281–1292. Available at: http://dx.doi.org/10.1016/j.cell.2014.08.011.

Gingold, H., Dahan, O. & Pilpel, Y., 2012. Dynamic changes in translational efficiency are deduced from codon usage of the transcriptome. Nucleic Acids Research, 40(20), pp.10053–10063. Available at: https://watermark.silverchair.com/gks772.pdf?token=AQECAHi208BE49Ooan9kkhW_Ercy7Dm3ZL_9Cf3qfKAc485ysgAAAcIwggG-BgkqhkiG9w0BBwagggGvMIIBqwIBADCCAaQGCSqGSIb3DQEHATAeBglghkgBZQMEAS4wEQQMEfTe9lBhk9unJKXhAgEQgIIBdezEM3fk7PHLJGZLYTb1_8Fq6YkXICElZuTB4mq6VLlEbHK3 [Accessed December 26, 2017].

Goodarzi, H. et al., 2016. Modulated expression of specific tRNAs drives gene expression and cancer progression. Cell, 165(6), pp.1416–1427. Available at: http://dx.doi.org/10.1016/j.cell.2016.05.046.

Hafner, A. et al., 2017. p53 pulses lead to distinct patterns of gene expression albeit similar DNA-binding dynamics. nature structural and molecular biology, 24(10), pp.840–847. Available at: https://www.nature.com/articles/nsmb.3452.pdf [Accessed June 26, 2019].

Hafner, A. et al., 2019. The multiple mechanisms that regulate p53 activity and cell fate. Nature Reviews Molecular Cell Biology, 20(4), pp.199–210. Available at: http://dx.doi.org/10.1038/s41580-019-0110-x.

Hanahan, D. & Weinberg, R.A., 2011. Hallmarks of cancer: The next generation. Cell, 144(5), pp.646–674.

Hanson, G. & Coller, J., 2017. Codon optimality, bias and usage in translation and mRNA decay. Nature Publishing Group, 19. Available at: https://www.nature.com/articles/nrm.2017.91.pdf [Accessed December 27, 2017].

Hernandez-Segura, A. et al., 2017. Unmasking Transcriptional Heterogeneity in Senescent Cells. Current Biology, 27(17), p.2652–2660.e4. Available at: https://linkinghub.elsevier.com/retrieve/pii/S0960982217308928 [Accessed June 26, 2019].

Hernandez-Alias, X. et al., 2020. Translational efficiency across healthy and tumor tissues is proliferation-related. Molecular Systems Biology, 16(3), pp.1–15.

Ho, T.-T. et al., 2015. Targeting non-coding RNAs with the CRISPR/Cas9 system in human cell lines. Nucleic acids research, 43(3), p.e17. Available at: http://www.pubmedcentral.nih.gov/articlerender.fcgi?artid=4330338&tool=pmcentrez&rendertype=abstract.

Kirchner, S. & Ignatova, Z., 2014. Emerging roles of tRNA in adaptive translation, signalling dynamics and disease. Nature Publishing Group, 16. Available at: www.nature.com/reviews/genetics [Accessed August 15, 2018].

Knight, J.R.P. et al., 2020. Control of translation elongation in health and disease. Disease Models & Mechanisms, 13(3), p.dmm043208.

Krylov, D.M. et al., 2003. Gene loss, protein sequence divergence, gene dispensability, expression level, and interactivity are correlated in eukaryotic evolution. Genome Research, 13(10), pp.2229–2235. Available at: www.genome.org [Accessed March 30, 2020].

Lowe, T.M. & Chan, P.P., 2016. tRNAscan-SE On-line: integrating search and context for analysis of transfer RNA genes. Nucleic Acids Research, 44. Available at: http://trna.ucsc.edu/tRNAscan-SE/. [Accessed August 30, 2018].

Milyavsky, M. et al., 2003. Prolonged Culture of Telomerase-Immortalized Human Fibroblasts Leads to a Premalignant Phenotype. Cancer Research, 63(21), pp.7147–7157.

Montague, T.G. et al., 2014. CHOPCHOP: a CRISPR/Cas9 and TALEN web tool for genome editing. Nucleic Acids Research, 42, pp.401–407. Available at: https://chopchop.rc.fas. [Accessed April 21, 2020].

Nagano, T. et al., 2016. Identification of cellular senescence-specific genes by comparative transcriptomics OPEN. Available at: www.nature.com/scientificreports [Accessed March 14, 2019].

Oki, T. et al., 2014. A novel cell-cycle-indicator, mVenus-p27K -, identifies quiescent cells and visualizes G0-G1 transition. Scientific Reports, 4(4012), pp.1–10.

Pan, T., 2018. Modifications and functional genomics of human transfer RNA. Cell Research, pp.1–10. Available at: https://www.nature.com/articles/s41422-018-0013-y.pdf [Accessed April 9, 2018].

Patil, A. et al., 2012. Cell Cycle Increased tRNA modification and gene-specific codon usage regulate cell cycle progression during the DNA damage response View supplementary material. Cell Cycle, 11, pp.3656–3665. Available at: https://www.tandfonline.com/action/journalInformation?journalCode=kccy20 [Accessed April 1, 2020].

Pavon-Eternod, M. et al., 2013. Overexpression of initiator methionine tRNA leads to global reprogramming of tRNA expression and increased proliferation in human epithelial cells. RNA, 19(4), p.461:466. Available at: http://rnajournal.cshlp.org/content/19/4/461.full.pdf [Accessed June 8, 2018].

Pavon-Eternod, M. et al., 2009. tRNA over-expression in breast cancer and functional consequences. Nucleic Acids Research, 37(21), pp.7268–7280.

Pérez-Mancera, P.A., Young, A.R.J. & Narita, M., 2014. Inside and out: The activities of senescence in cancer. Nature Reviews Cancer, 14(8), pp.547–558.

Perucca, P. et al., 2009. Loss of p21CDKN1A impairs entry to quiescence and activates a DNA damage response in normal fibroblasts induced to quiescence. Cell Cycle, 8(1), pp.105–114.

Pinello, L. et al., 2016. Analyzing CRISPR genome-editing experiments with CRISPResso. Nature Biotechnology, 34(7), pp.695–697.

Presnyak, V. et al., 2015. Codon Optimality Is a Major Determinant of mRNA Stability. Cell, 160, pp.1111–1124. Available at: http://dx.doi.org/10.1016/j.cell.2015.02.029http://dx.doi.org/10.1016/j.cell.2015.02.029 [Accessed July 9, 2018].

Rak, R., Dahan, O. & Pilpel, Y., 2018. Repertoires of tRNAs: The Couplers of Genomics and Proteomics. Annual Review of Cell and Developmental Biology Annu. Rev. Cell Dev. Biol, 34, pp.20–21. Available at: https://doi.org/10.1146/annurev-cellbio-100617- [Accessed August 28, 2018].

Rapino, F. et al., 2018. Codon-specific translation reprogramming promotes resistance to targeted therapy. Natu, 558, pp.605–609. Available at: https://www.nature.com/articles/s41586-018-0243-7.pdf [Accessed July 5, 2018].

Dos Reis, M., Savva, R. & Wernisch, L., 2004. Solving the riddle of codon usage preferences: a test for translational selection. Nucleic Acids Research, 32(17), pp.5036–5044. Available at: https://watermark.silverchair.com/gkh834.pdf?token=AQECAHi208BE49Ooan9kkhW_Ercy7Dm3ZL_9Cf3qfKAc485ysgAAAcEwggG9BgkqhkiG9w0BBwagggGuMIIBqgIBADCCAaMGCSqGSIb3DQEHATAeBglghkgBZQMEAS4wEQQMwXyspAuNa7jKbuwKAgEQgIIBdE1Hq2y3SdnpvoS1iEg_PnopsTFKtk4hiI1vsEjb731gE0MM [Accessed February 15, 2018].

Rew, D.A. & Wilson, G.D., 2000. Cell production rates in human tissues and tumours and their significance. Part II: clinical data. European Journal of Surgical Oncology (EJSO), 26(4), pp.405–417. Available at: https://linkinghub.elsevier.com/retrieve/pii/S0748798399909071 [Accessed August 20, 2019].

Rufini, A. et al., 2013. Senescence and aging: the critical roles of p53. Oncogene, 32, pp.5129–5143.

Ruijtenberg, S. & Van Den Heuvel, S., 2016. Coordinating cell proliferation and differentiation: Antagonism between cell cycle regulators and cell type-specific gene expression. Cell Cycle, 15(2), pp.196–212. Available at: http://www.tandfonline.com/doi/pdf/10.1080/15384101.2015.1120925?needAccess=true [Accessed December 5, 2017].

Sambrook, J. & Russell, D.W., 2001. Molecular Cloning N. Irwin, ed., Cold Spring Harbor Laboratory Press.

Santos, M. et al., 2019. tRNA Deregulation and Its Consequences in Cancer. Trends in Molecular Medicine. Available at: https://linkinghub.elsevier.com/retrieve/pii/S1471491419301285 [Accessed July 2, 2019].

Shim, G. et al., 2017. Therapeutic gene editing: delivery and regulatory perspectives. Nature Publishing Group, 38, pp.738–753. Available at: www.nature.com/aps [Accessed April 3, 2020].

Sosa, M.S., Bragado, P. & Aguirre-Ghiso, J.A., 2014. Mechanisms of disseminated cancer cell dormancy: An awakening field. Nature Reviews Cancer, 14(9), pp.611–622.

Spencer, S.L. et al., 2013. The Proliferation-Quiescence Decision Is Controlled by a Bifurcation in CDK2 Activity at Mitotic Exit. Cell, 155, pp.369–383. Available at: http://dx.doi.org/10.1016/j.cell.2013.08.062. [Accessed December 4, 2019].

Thornlow, B. et al., 2018. Transfer RNA genes experience exceptionally elevated mutation rates. PNAS, p.229906. Available at: https://www.biorxiv.org/content/early/2018/06/05/229906 [Accessed August 28, 2018].

Tong, S. et al., 2019. Engineered materials for in vivo delivery of genome-editing machinery. Nature Reviews Materials, 4(11), pp.726–737. Available at: http://dx.doi.org/10.1038/s41578-019-0145-9.

Wesley Overton, K. et al., 2014. Basal p21 controls population heterogeneity in cycling and quiescent cell cycle states. Proceedings of the National Academy of Sciences of the United States of America, 111(41), pp.E4386–E4393.

Yao, G., 2014. Modelling mammalian cellular quiescence. Interface Focus, 4.

Yuen, G. et al., 2017. CRISPR/Cas9-mediated gene knockout is insensitive to target copy number but is dependent on guide RNA potency and Cas9/sgRNA threshold expression level. Nucleic acids research, 45(20), pp.12039–12053. Available at: http://dx.doi.org/10.1093/nar/gkx843.

Zhang, Z. et al., 2018. Global analysis of tRNA and translation factor expression reveals a dynamic landscape of translational regulation in human cancers. Communications Biology, 1(234). Available at: https://www.nature.com/articles/s42003-018-0239-8.pdf [Accessed June 18, 2019].

Zheng, G. et al., 2015. Efficient and quantitative high-throughput tRNA sequencing. Nature Methods.

